# Extrinsic KRAS signaling shapes the pancreatic microenvironment through fibroblast reprogramming

**DOI:** 10.1101/2021.08.26.457660

**Authors:** Ashley Velez-Delgado, Katelyn L. Donahue, Kristee L. Brown, Wenting Du, Valerie Irizarry-Negron, Rosa E. Menjivar, Emily L. Lasse Opsahl, Nina G. Steele, Stephanie The, Jenny Lazarus, Veerin R. Sirihorachai, Wei Yan, Samantha B. Kemp, Samuel A. Kerk, Murali Bollampally, Fatima Lima, Costas A. Lyssiotis, Arvind Rao, Howard C. Crawford, Filip Bednar, Timothy L. Frankel, Yaqing Zhang, Marina Pasca di Magliano

**Affiliations:** Department of Cell and Developmental Biology, University of Michigan, Ann Arbor, MI 48109 USA; Cancer Biology Program, University of Michigan, Ann Arbor, MI 48109 USA; Department of Surgery, University of Michigan, Ann Arbor, MI 48109 USA; Cellular and Molecular Biology Program, University of Michigan, Ann Arbor, MI 48109 USA; Department of Computational Medicine and Bioinformatics, University of Michigan, Ann Arbor, MI 48109 USA; Molecular and Cellular Pathology Program, University of Michigan, Ann Arbor, MI 48109 USA; Life Sciences and Arts college, University of Michigan, Ann Arbor, MI 48109 USA; Department of Molecular and Integrative Physiology, University of Michigan, Ann Arbor, MI 48109 USA; Rogel Cancer Center, University of Michigan, Ann Arbor, MI 48109 USA; Department of Internal Medicine, Division of Gastroenterology and Hepatology, University of Michigan, Ann Arbor, MI 48109 USA; Michigan Institute of Data Science (MIDAS), University of Michigan, Ann Arbor, MI 48109 USA; Department of Radiation Oncology, University of Michigan, Ann Arbor, MI 48109 USA

**Keywords:** pancreatic cancer, KRAS, transformation, fibroblasts, macrophages, cellular plasticity

## Abstract

Oncogenic KRAS is the hallmark mutation of human pancreatic cancer and a driver of tumorigenesis in genetically engineered mouse models of the disease. While the tumor cell-intrinsic effects of oncogenic Kras expression have been widely studied, its role in regulating the extensive pancreatic tumor microenvironment is less understood. Using a genetically engineered mouse model of inducible and reversible oncogenic Kras expression and a combination of approaches that include mass cytometry and single cell RNA sequencing, we have discovered that non-cell autonomous (i.e., extrinsic) oncogenic KRAS signaling reprograms pancreatic fibroblasts, activating an inflammatory gene expression program. As a result, fibroblasts become a hub of extracellular signaling, mediating the polarization and function of pro-tumorigenic myeloid cells while also preventing tissue repair. Our study provides fundamental new knowledge on the mechanisms underlying the formation of the fibroinflammatory stroma in pancreatic cancer and highlights stromal pathways with the potential to be exploited therapeutically.

## INTRODUCTION

Pancreatic ductal adenocarcinoma (PDA) is the third leading cause of cancer-related death in the United States and is counted among the most lethal malignancies, with an expected 5-year survival of about 10% (1). Over 90% of PDA instances harbor an oncogenic mutation in the *KRAS* gene, most commonly *KRAS^G12D^* (2, 3). Autopsy studies have revealed that over 75% of the population harbors pre-neoplastic lesions in the pancreas linked to mutations in *KRAS*, yet pancreatic cancer is relatively rare (4, 5). This finding is reproduced in mouse models of the disease, where widespread epithelial expression of oncogenic Kras leads to cancer with long latency. Why some KRAS-mutant lesions maintain an indolent pre-neoplastic state while others progress to deadly invasive cancer is a fundamental gap in knowledge.

A longstanding question has been the identity of the cell of origin to pancreatic cancer. The most frequently utilized genetically engineered mouse models of pancreatic cancer express Kras^G12D^ broadly across the pancreas epithelium upon Cre recombination. Two of the most common Cre drivers include the Pdx1 promoter (Pdx1-Cre;Kras^LSL-G12D/+^) and insertion into the Ptf1a locus (Ptf1a^Cre/+^;Kras^LSL-G12D/+^); both of these models are commonly referred to as KC (6) (7). Different precursor lesions to pancreatic cancer have been described in human patients (8}), of which Pancreatic Intraepithelial Neoplasia (PanIN) is the most common. KC mice undergo a stepwise carcinogenesis process that mimics the progression of human disease, including PanIN formation (6) (7). While in mouse models both acinar cells and ductal cells can give rise to pancreatic cancer, acinar cell origin is the most frequent for both PanIN and cancer (9), (10), (11), (12).

Acinar cells are highly plastic – upon tissue damage, they downregulate expression of digestive enzymes and acquire a duct-like differentiation status through a process known as acinar-to-ductal metaplasia (ADM). Acinar cells that have undergone ADM can be distinguished from ductal cells due to their unique expression of acinar progenitor factors (13). In the context of acute injury, ADM is reversible, and the acinar parenchyma is re-established over time. However, expression of oncogenic Kras prevents the recovery of ADM and the duct-like cells instead undergo neoplastic transformation (14). The cell intrinsic mechanisms regulating the balance between cellular plasticity and carcinogenesis have been extensively studied. ADM is driven by fundamental changes in the transcriptional regulation of acinar cells, with acinar transcription factors restraining and ductal transcription factors promoting this process (for review see (15)). For example, reduction of expression of the acinar transcription factors Ptf1a, Mist1 or Nr5a2 promote transformation, while forced expression of Mist 1 and Ptf1a protect acinar cells from de-differentiation (16–21). ADM is also regulated by intracellular signaling, including a requirement for epithelial MAPK activation (22, 23). Recently, epigenetic reprogramming driven by oncogenic Kras has emerged as a key determinant of progression/redifferentiation of acinar cells upon injury (24).

While the cell autonomous (i.e., intrinsic) effects of oncogenic Kras activation have been extensively studied, the non-cell autonomous effects are less understood. Here, we set out to investigate how oncogenic Kras-expressing cells affect the microenvironment, which we refer to as “extrinsic” Kras signaling. The transdifferentiation of acinar cells and formation of ADM/PanIN are accompanied by changes in the surrounding microenvironment, with activation of fibroblasts (for review see Helms, 2020, 32014869) and infiltration of immune cells. The latter are largely immune suppressive (25), and include myeloid cells that both suppress T cell responses and directly promote pancreatic cancer progression (26, 27). Myeloid cells support ADM (28) and are required for sustaining ADM and promoting progression to PanIN (29). Conversely, myeloid cells are also required for tissue repair both in the setting of injury and redifferentiation (29, 30). Further, experimental induction of acute pancreatitis with its associated inflammation synergizes with oncogenic Kras to accelerate the formation of ADM and PanIN (31, 32). Thus, inflammation both accompanies and promotes carcinogenesis, but the relationship between epithelial cells and surrounding stroma in this process remains unclear.

In this study, we investigate how extrinsic signaling mediated by oncogenic Kras regulates the formation and maintenance of the fibroinflammatory stroma and the cellular crosstalk during the initiation of pancreatic cancer. To this end, we have used a genetically engineered mouse model of inducible and reversible oncogenic Kras expression previously described by our group and others (33, 34). By exploiting our ability to activate and inactivate oncogenic Kras expression at will, we have dissected the events immediately following the inception of oncogenic Kras expression and observed how continuous oncogenic Kras signaling both maintains acinar transdifferentiation and directs the surrounding microenvironment. We have discovered that fibroblast reprogramming occurs during the earliest stages of carcinogenesis and in turn drives the tumor-promoting functional status of myeloid cells infiltrating the pancreas. Consequently, approaches to reprogram fibroblasts during carcinogenesis should be explored to prevent, and possibly reverse, KRAS-driven carcinogenesis.

## RESULTS

### Extrinsic signaling by oncogenic Kras reprograms the pancreas microenvironment

To study the recruitment of immune cells by oncogenic Kras expressing-tumor cells, we used Ptf1a^Cre/+^;R26^rtTa-ires-EGFP/rtTa-ires-EGFP^;TetO-Kras^G12D^ mice (hereby **iKras\***) that expresses oncogenic Kras^G12D^ (hereby Kras*) in pancreatic epithelial cells in an inducible and reversible manner upon Doxycycline (DOX) administration (33). Initially, we activated Kras* in adult mice (6-12 weeks old) by feeding iKras* mice or littermate controls (lacking either Cre or Kras* expression) DOX chow for 3 days, 1 week or 2 weeks (See scheme in **Fig. 1A**). Pancreata from iKras* mice appeared largely histopathologically normal 3 days following Kras* activation, with the parenchyma largely composed of acinar cells and CK19 expression confined to the ducts (**Fig. 1B** and **C** and quantification of ADM in **Fig. S1A**). One week after Kras* induction, we observed focal ADM accompanied by an increase in the proliferation marker Ki67 that became more widespread by 2 weeks (**Fig. 1B** and **Fig. S1B**). At the 3-day timepoint, we observed patchy epithelial expression of phosphorylated ERK (p-ERK), indicating activation of the MAPK pathway (**Fig. 1D**) in otherwise histologically normal areas. At later time points, areas of apparent ADM had elevated p-ERK staining (**Fig. 1D**). The limited expression of p-ERK even as all pancreas epithelium express Kras* is consistent with the known requirement of stimulation by various upstream factors to fully activate Kras* signaling, including myeloid-derived EGFR ligands (35)) (36, 37).

**Figure 1.**
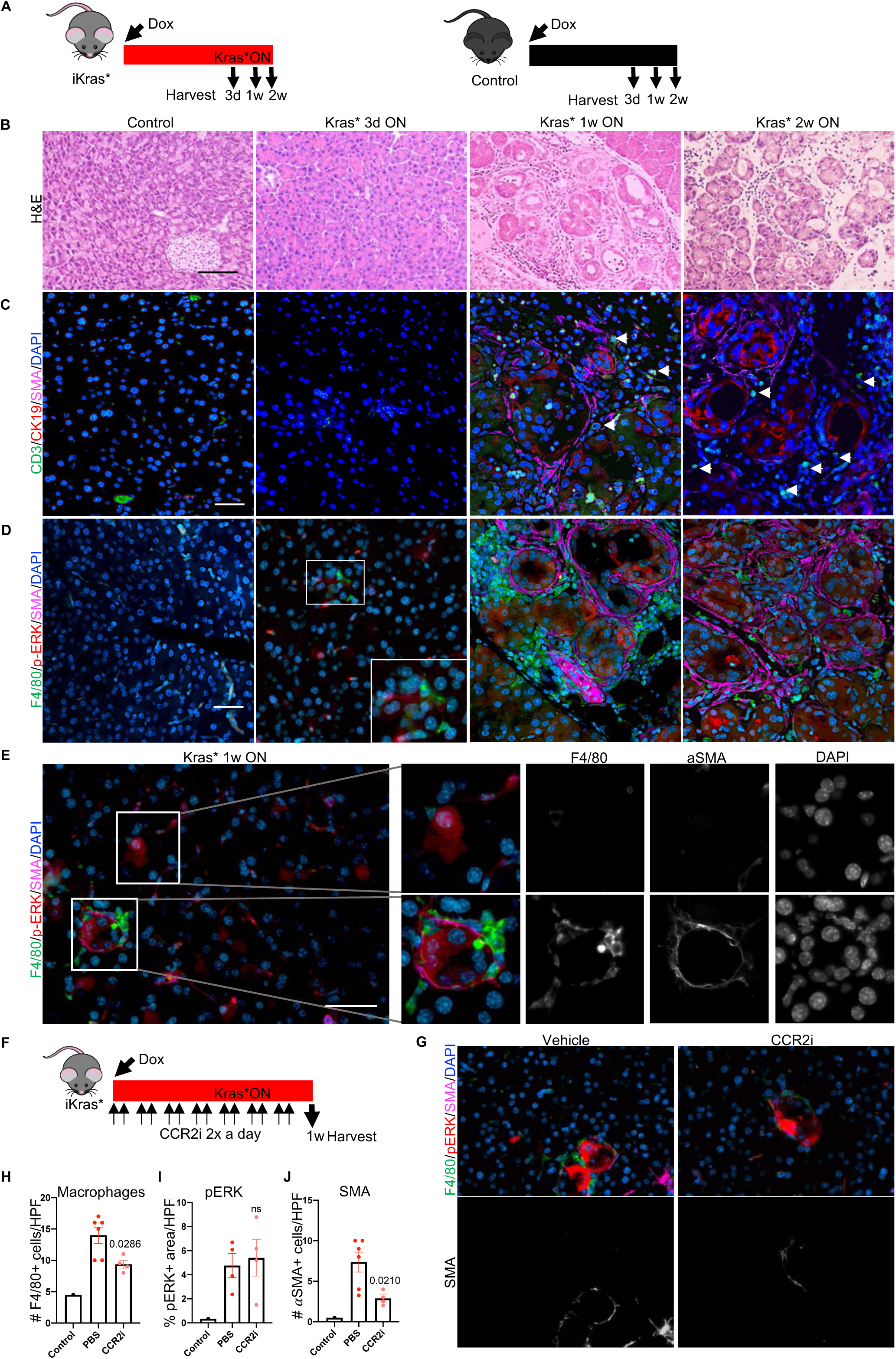
*Oncogenic Kras drives the recruitment of myeloid cells prior to acinar transdifferentiation.* **A**, Experimental design. Control (lacking either the Kras or Ptf1a^Cre^ allele) and iKras* mice were given Doxycycline (DOX) chow to activate oncogenic Kras (Kras*) in the iKras* mice and pancreata were harvested either at 3 days (3d), 1 week (1w) or 2 weeks (2w). N=3-5 mice per group. **B**, Representative images of Hematoxylin and Eosin (H&E) staining of control and iKras* pancreata at the indicated time points. N=3-5 mice per group. Scale bar 100μm. **C**, Representative images of CD3 (green), CK19 (red) and SMA (magenta) co-immunofluorescent staining in control and iKras* pancreata at the indicated time points, arrows pointing to CD3^+^ T cells. N=3 mice per group. Scale bar 50μm. **D,** Representative images of F4/80 (green), p- ERK (red) and SMA (magenta) co-immunofluorescent staining in control and iKras* pancreata at the indicated time points. N=3 mice per group. Scale bar 50μm. **E,** Representative images of F4/80 (green), p-ERK (red) and SMA (magenta) co-immunofluorescence staining of an iKras* pancreas after activating Kras* for 1 week. Regions with high expression of p-ERK are enlarged and single channel images are included. N=3 mice. Scale bar 50μm. **F,** Experimental design. iKras* mice were given DOX to turn ON Kras* and were treated at the same time with a CCR2 inhibitor (CCR2i) or PBS twice a day for a week. N=6 mice per group. **G,** Representative images of F4/80 (green), p-ERK (red) and SMA (magenta) co-immunofluorescent staining. Bottom row: single channel images for SMA. Scale bar 50μm. Quantification of F4/80^+^ macrophages **(H)**, p-ERK^+^ cells **(I)** and SMA^+^ activated fibroblasts **(J)** for the experiment described in F. Data is measured as number of cells per high power field (HPF) (400X for F4/80 and SMA, 200X for p-ERK), and expressed as mean ± SEM. The statistic difference between PBS and CCR2i was determined by two-tailed Mann-Whitney or unpaired t-test. See also Figure S1

Next, we investigated the dynamics of immune infiltration and the presence of activated fibroblasts (as marked by expression of smooth muscle actin, SMA). Co-immunofluorescence and immunohistochemistry analysis revealed the presence of macrophages surrounding untransformed p-ERK-expressing acinar cells as early as 3 days post-Kras* activation (**Fig. 1D** and **S1C**), while T cells were rare at this stage (**Fig. 1C and S1D**). Macrophages increased by 1 and 2 weeks post-Kras* activation, and T cells were easily detected at these timepoints as well **(Fig. 1C, S1C and 1D)**. Areas of ADM at this stage were also surrounded by SMA^+^ fibroblasts (**Fig. 1C and 1D**). We then examined the areas of the pancreas that were morphologically normal at the 1-week timepoint (**Fig. 1E**). Interestingly, high-magnification images revealed morphologically normal clusters of acini with elevated p-ERK surrounded by both macrophages and a thin layer of SMA^+^ fibroblasts, indicating that changes in the microenvironment precede acinar transdifferentiation. Careful analysis of the tissue from the 3-day timepoint also revealed some p-ERK- expressing acini with surrounding macrophages but without SMA^+^ fibroblasts (**Fig. 1D**). When we evaluated neutrophil infiltration (Ly6b^+^) at this timepoint, we observed them more frequently in ADM than in histologically normal areas, suggesting that their recruitment is later in the transdifferentiation process than macrophages (**Fig. S1E**). Taken together, our findings are consistent with a model whereby macrophage infiltration and p-ERK activation occur in tandem, followed by fibroblast activation and ultimately acinar transdifferentiation.

While the pancreas is home to resident macrophages that expand during carcinogenesis (38), CCR2^+^ monocytes are also recruited to the pancreas and give rise to macrophages (39, 40). To understand the contribution of monocyte-derived macrophages to the establishment of ADM, we treated iKras* mice with the CCR2 inhibitor, PF-04178903, (hereby CCR2i) (41) or vehicle control, concurrently with Kras* activation (**Fig. 1F**). Tissue was harvested 1 week following treatment. ADM was rare in both cohorts, and we observed no difference in its prevalence (**Fig. S1F**). There was an ∼1/3 decrease in the number of macrophages in the tissue of CCR2i-treated pancreata compared to vehicle controls, consistent with a model by which recruitment of monocyte-derived macrophages had been prevented, while tissue resident macrophages, known to be prevalent at this stage (38), remained unaffected (**Fig. 1G and 1H**). We observed no difference in the levels of p-ERK staining, indicating that infiltrating macrophages are not required to initiate MAPK signaling at this stage (**Fig. 1G** and **1I**), possibly as resident macrophages serve as a sufficient source of EGFR ligands (35). Interestingly, we observed a reduction in the number of activated fibroblasts surrounding p-ERK expressing acinar clusters (**Fig. 1G** and **1J**). These data suggest that monocyte-derived macrophages are recruited through the CCL2/CCR2 axis following epithelial activation of oncogenic Kras and are required for fibroblast activation during Kras*-induced acinar transdifferentiation.

### Epithelial oncogenic Kras* is required to maintain acinar transdifferentiation and fibroblast accumulation

Acute pancreatitis synergizes with oncogenic Kras to drive neoplastic transformation (31–33). Accordingly, we placed iKras* and control mice on DOX and then induced acute pancreatitis (see scheme in **Fig. 2A**). Pancreata in iKras* mice presented with widespread ADM and low grade PanINs (with intracellular mucin accumulation as shown by PAS staining) and accumulation of fibrotic stroma (**Fig. 2B and 2C**). In contrast, control mice treated in parallel had normal pancreas architecture. iKras* pancreata were also marked by extensive collagen deposition, fibroblast expansion and activation (as detected by podoplanin and SMA, respectively) and epithelial p-ERK accumulation (**Fig. 2C, 2D** and **2E**). To study the effect of Kras* inactivation on the surrounding microenvironment, we replaced DOX chow with DOX-free chow to inactivate Kras* expression in iKras* mice (for 3 days or 1 week) 3 weeks post-administration of acute caerulein (see scheme in **Fig. 2A**). The 3-day and 1-week timepoints were chosen to coincide with the early and mid-phases of the pancreatic remodeling process, as we have previously shown (33). As previously described, Kras* inactivation led to a drastic reduction in p-ERK and redifferentiation of acinar cells progressively over time (**Fig. 2B, 2E**) (29). We then investigated changes in the stroma following inactivation of Kras*. We observed a reduction in both total fibroblasts (podoplanin^+^ cells) and of activated, SMA^+^ fibroblasts, as well as a reduction in collagen deposition (**Fig. 2C, 2D** and **2E** and quantification in **2F and 2G**). These results are consistent with a requirement for continued oncogenic Kras* activity to maintain fibroblast expansion and pancreatic fibrosis.

**Figure 2.**
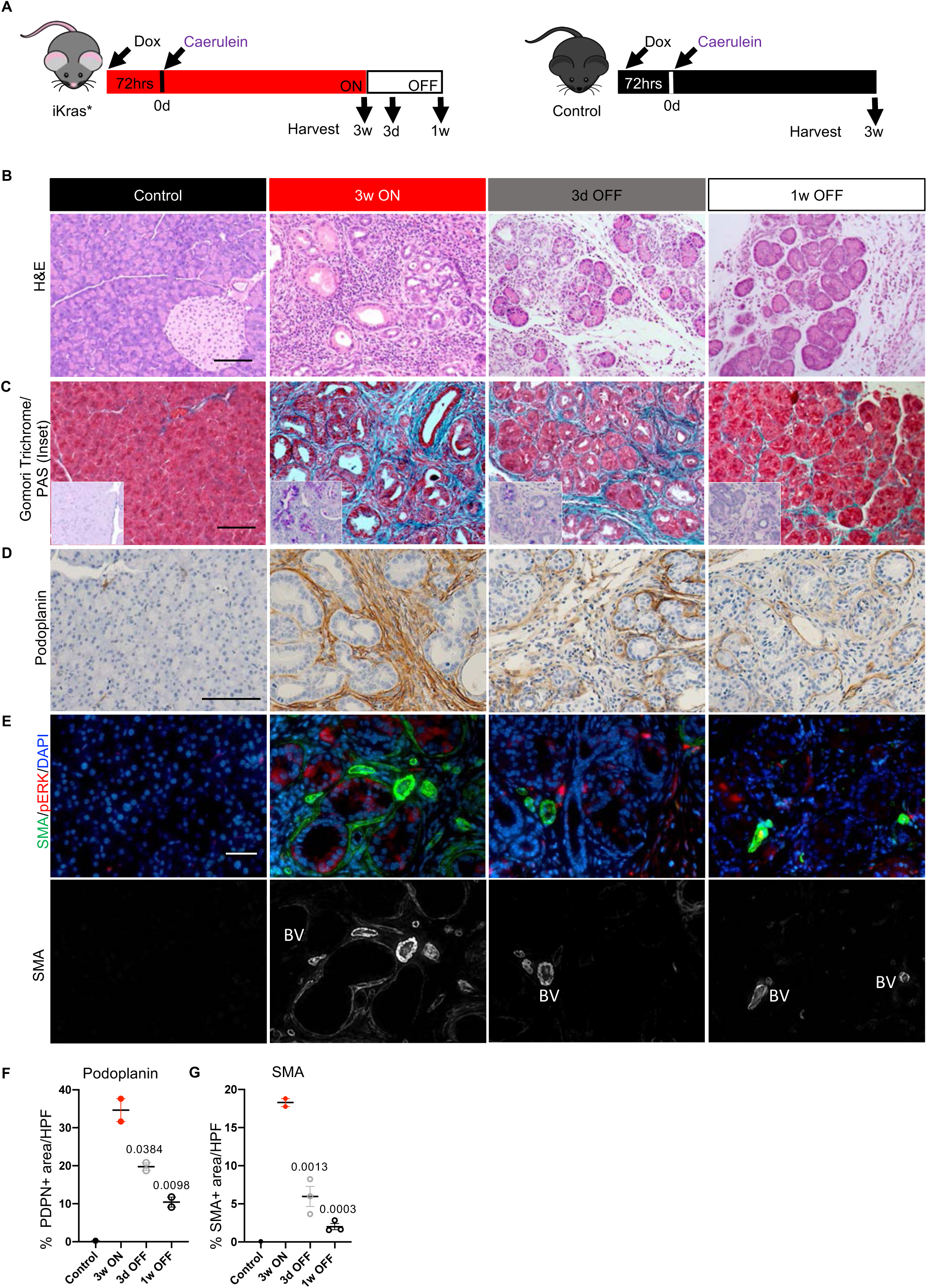
*Sustained expression of Kras is required to maintain PanIN and fibrosis.* **A,** Experimental design. Wild type control and iKras* mice were given DOX chow to activate Kras* followed by induction of acute pancreatitis. Mice were then left on DOX chow for three weeks, at which point pancreata were harvested (3w ON) or the DOX chow was removed and the pancreata were harvested after 3 days or 1 week (labeled 3d OFF or 1w OFF respectively). N=8 mice per group. **B,** Representative images of H&E staining of control and iKras* pancreata at the indicated time points, N=8 mice per group. Scale bar 100μm. **C,** Representative images of Gomori Trichrome and Periodic Acid-Schiff Stain (PAS; inset) of control or iKras* pancreata at the indicated time points, N=2-3 mice per group. Scale bar 100μm. **D,** Immunostaining for Podoplanin (Scale bar 100μm). **E,** Immunofluorescent staining for SMA (green) and p-ERK (red). Scale bar 50μm. N=2-3 mice per group. Quantification of Podoplanin **(F)** and SMA staining **(G)**, shown as mean ± SEM. Statistical differences were determined by multiple ANOVA, all compared to 3w ON group.

### Extrinsic signaling from Kras* transformed cells drives myeloid polarization

Myeloid cells infiltrate early during pancreatic carcinogenesis and are required for sustained ADM and progression to PanIN (28,29,42). Therefore, we endeavored to study the effect of Kras* inactivation on myeloid cell infiltration in the pancreas. For this purpose, we placed iKras* and control mice on DOX and then induced acute pancreatitis (See scheme in **Fig. 3A**). Three weeks later, animals were randomized to DOX or DOX-free chow to inactivate Kras* expression for 3 days or 1 week. We then performed a combination of flow cytometry with an antibody panel enriched for myeloid cell markers (Table S1; gating strategy in **Fig. S2A**) and immunostaining, including multiplex immunofluorescent staining (Opal multiplex IHC, Akoya). As expected, we observed an increase in CD45^+^ immune cells in the iKras* mice at 3w ON compared to control mice (**Fig. 3B and S2C**). This increase included total myeloid cells (CD45^+^CD11b^+^) as well as macrophages (CD45^+^CD11b^+^F4/80^+^) and immature myeloid cells (CD45^+^CD11b^+^F4/80^-^ Ly6C^+^Ly6G^+^, often referred to as MDSCs) (**Fig. 3B, S2B and S2C**), consistent with the increase in leukocyte infiltration observed in KC mice (25). Surprisingly, Kras* inactivation resulted in little to no reduction in total immune infiltration, total myeloid cells, or macrophages (**Fig. 3B, S2B and S2C**). As myeloid cell numbers did not change upon Kras* inactivation, we sought to determine whether their polarization status was affected. For this purpose, we used mass cytometry (CyTOF) which allowed us to use a panel of ∼20 antibodies concurrently (see Table S2). We limited the scope of the CyTOF experiment to the 3 weeks Kras* ON (N=2) and 3 days OFF (N=3) timepoints in order to highlight the short-term effect of Kras* inactivation, prior to extensive tissue remodeling. We visualized the multi-dimensional CyTOF data using tSNE plots (t-distributed stochastics neighbor embedding) and identified 16 distinct immune cell clusters (**Fig. 3C** and **S2D**). We observed no significant change in the T cell clusters upon Kras* inactivation, although there was a trend towards an increase in CD4^+^ T cells (**Fig. S2E**). The myeloid immune landscape was complex, with multiple populations, including heterogeneous macrophage subtypes (**Fig. 3C, 3D** and **S2D**). While some populations did not change or tended towards an increase when we compared Kras* ON to Kras* OFF samples, (such as iNOS^+^ macrophages, **Fig. S2F**), other populations drastically decreased. Notably, CCR1^+^CD206^+^ macrophages, immunosuppressive ARG1^+^ monocytic macrophages and MDSCs, defined as CD45^+^CD11b^+^F4/80^-^Ly6C^+^Ly6G^+^, decreased upon Kras* inactivation (**Fig. 3D**). The decrease in macrophages and ARG1^+^ macrophages was also validated by co-immunofluorescent and multiplex immunofluorescent staining (**Fig. 3E, S3A, S3B** and **S3C**). Furthermore, JAK/STAT3 signaling, known to promote the immune-suppressive function of macrophages, (43), is similarly regulated by epithelial oncogenic Kras (**Fig. 3F**).

**Figure 3.**
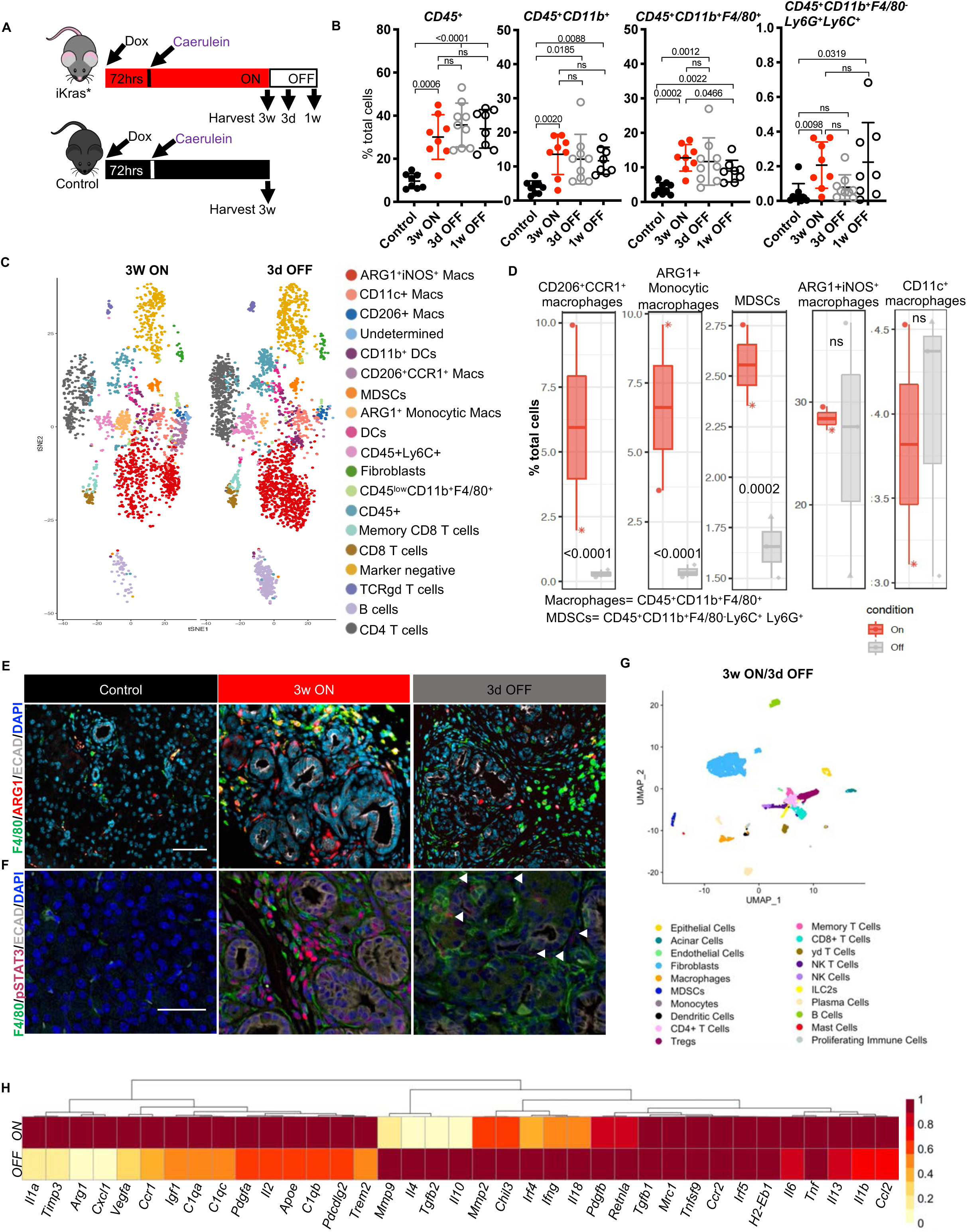
*Oncogenic Kras regulates myeloid cell polarization status.* **A,** Experimental Design. N=8-9 mice per group. **B,** Quantification of flow cytometry results. Immune cell populations are expressed as percentage of total cells in control or iKras* pancreata at the indicated time points, shown as mean ± SD. Statistical analysis by multiple comparison ANOVA and multiple comparison Kruskal Wallis. **C,** CyTOF analysis of iKras* pancreata visualized by tSNE plot for 3w ON and 3d OFF timepoints. Nineteen distinct cell clusters were identified by FLowSOM using 18 markers and 1061 randomly selected cells per group. N=2-3 mice per group. **D,** Quantification of specific cell populations, as indicated. The statistical analysis by Wrapper function. **E,** Immunostaining for F4/80 (green), Arg1 (red), and Ecad (White) in control and iKras* pancreata at the indicated time points. N=3 mice per group. Scale bar 50μm. **F,** Immunostianing for of F4/80 (green), pSTAT3 (Magenta), and Ecad (White). N=3 mice per group. Scale bar 50μm. **G,** Uniform and manifold approximation and projection (UMAP) visualization of single-cell RNA-sequencing data showing unsupervised clustering of cells from combined iKras* pancreatic samples (3w ON, N=2 and 3d OFF, N=3). Each color represents a distinct cellular cluster. **H,** Heatmap showing the averaged single-cell RNA sequencing expression data (relative to the highest expressor) for genes in macrophages selected from a curated list of macrophage polarization and functional markers. See also Figure S2, Figure S3

As CyTOF and immunostaining are limited by available antibodies, we performed single cell RNA sequencing (scRNAseq) to fully elucidate how epithelial Kras* expression alters the microenvironment, with special interest in infiltrating myeloid cells. We harvested pancreata from mice with Kras* ON for 3 weeks plus 3 days (N=2, pooled for submission) or ON for 3 weeks and then OFF for 3 days (N=3, pooled for submission). The experiment was designed so that pancreata were harvested and processed at the same time to avoid batch effects. The multidimensional scRNAseq data included a total of 5,073 cells overall, with 1,984 cells from the 3w ON sample and 3,089 cells from the 3d OFF sample. The data was analyzed using the Seurat package in R (version 3.2.2) and visualized by UMAP (uniform and manifold approximation and projection) (**Fig. 3G**). By comparing the transcriptome of each cluster to known markers of epithelial, immune, and stromal cell types (**Fig. S4A**), we were able to identify acinar cells, CK19^+^ epithelial cells, fibroblasts, endothelial cells, and multiple populations of CD45^+^ immune cells. Similar varieties of cells were present in both the ON and OFF condition, allowing us to compare gene expression changes upon inactivation of Kras* within each cell population. We first assessed gene expression changes in the macrophage population and observed changes in polarization markers (**Fig. 3H**). Notably, tumor-associated macrophage (TAM) markers such as *Arg1* and *Ccr1* were downregulated upon Kras* inactivation. We also observed downregulation of Apolipoprotein E (*Apoe*) – a secreted protein that our group recently described as highly expressed in TAMs and as promoting immune suppression in invasive pancreatic cancer (44). Additionally, we observed downregulation of the complement genes *C1qa*, *C1qb, C1qc* and the transcription factor *Trem2*, which together define TAMs in several malignancies (45). Our group has also recently described a population of *C1qa*^+^*C1qb*^+^*Trem2*^+^ macrophages enriched in human and mouse pancreatic cancer both at the primary tumor and in liver metastases (46). These findings are consistent with TAM gene expression markers being activated in macrophages during the onset of neoplasia. Interestingly, inactivation of Kras* induces a “repair-associated” gene expression program, including *Tgfb, Il10*, *Il4*, *Pdgfb* and *Mmps* (**Fig. 3H**). Together, we show that oncogenic Kras inactivation does not alter overall immune infiltration, but drastically reprograms macrophage gene expression.

### Epithelial Kras* drives inflammatory reprogramming of pancreatic fibroblasts

We next sought to understand whether modulation of Kras* expression in epithelial cells influenced extracellular crosstalk in such a way as to account for our observed changes in macrophage reprogramming. To achieve this, we averaged the expression levels of genes from each cell population and looked for expression of known ligand and receptor pairs using a published list further curated in our laboratory (47, 48). Pairing of ligand and receptor-expressing populations gave us a list of potential predicted interactions within the tissue (**Fig. 4A**). We then plotted interactions that were significantly downregulated upon Kras* inactivation. Strikingly, the vast majority of Kras*-driven interactions connected fibroblast ligands with receptors on epithelial cells and a multitude of immune cells (**Fig. 4B, 4C**). These data suggest that fibroblasts are reprogrammed when exposed to Kras*-expressing epithelial cells, becoming a key signaling hub driving changes in the immune microenvironment. We identified multiple cytokines and chemokines expressed in fibroblasts in an epithelial Kras*-dependent manner **(Fig. 4D)**; among those, we selected the genes that were mainly expressed by fibroblasts, including *Il33, Il6, Cxcl1* and the inflammatory mediators *Saa1* and *Saa3* (the ortholog of human *SAA1*). While expression of these factors was not confined to fibroblasts, it was highest in this cell population (**Fig. 4E** and **S4B**). Conversely, the respective receptors for these secreted factors were expressed by immune cells, including *Cxcr2* (CXCL1 receptor) on immature myeloid cells, *Il6r* (IL6 receptor)*, P2rx7 and Scarb1* (SAA3 and SAA1 receptors) on several myeloid populations, and *Il1rl1* (IL33 receptor also known as ST2) on macrophages, Tregs, mast cells, and innate lymphoid cells type 2 (ILC2). Thus, fibroblasts extrinsically reprogrammed by Kras* epithelial cells express inflammatory molecules that bind receptors on myeloid cells, in turn activating their canonical downstream signaling pathways including JAK/STAT3 signaling. Accordingly, p-STAT3, a readout of active JAK/STAT3 signaling, is elevated in macrophages infiltrating PanIN-bearing pancreas (**Fig. 3F**).

**Figure 4.**
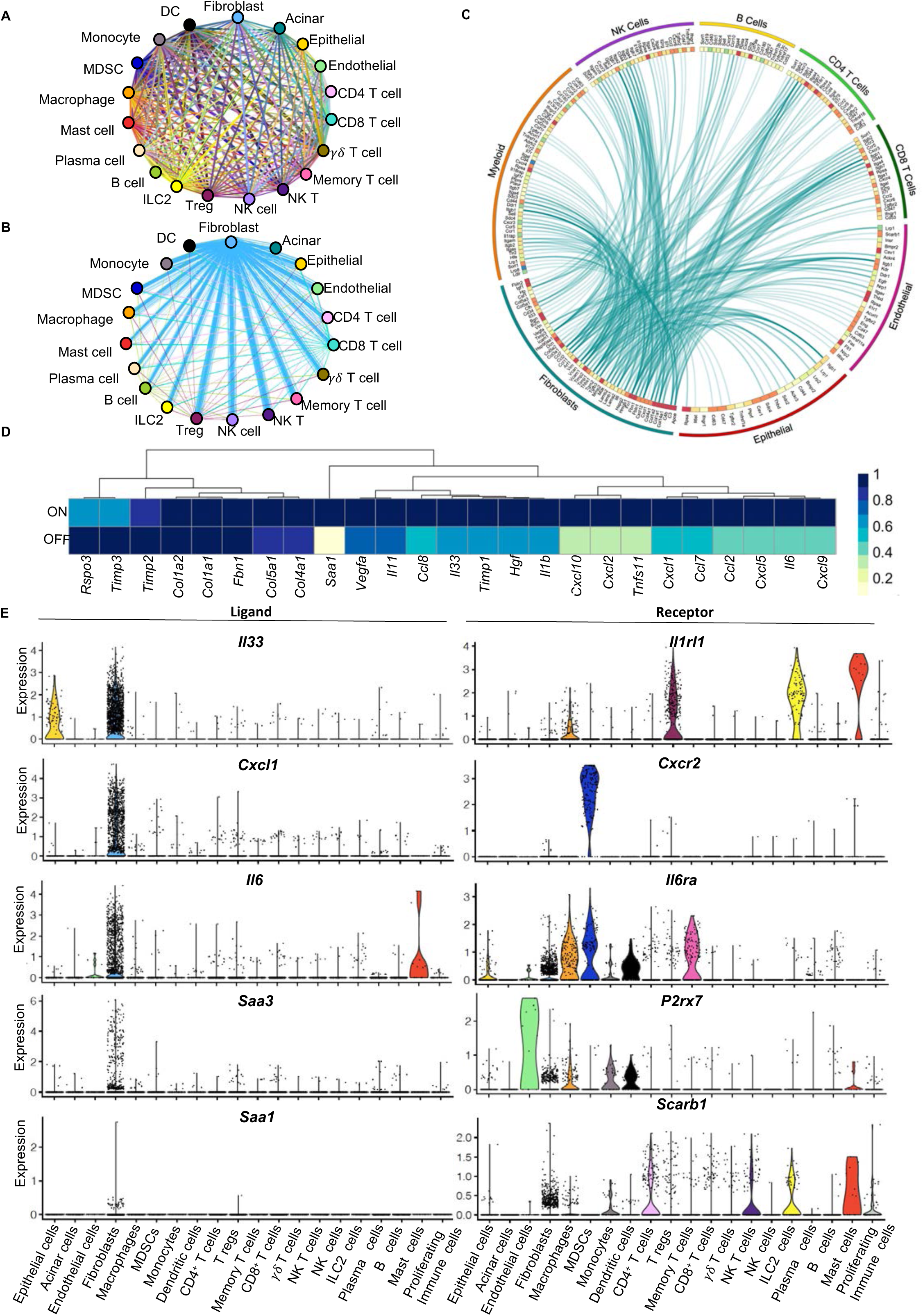
*Fibroblasts are important mediators of the immune infiltration.* **A,** Interactome analysis showing all predicted ligand-receptor interactions between different cell populations identified in iKras* pancreatic single-cell RNA sequencing analysis. Each line represents a ligand-receptor pair, color-coded by the cell-type expressing the ligand. **B,** Differential interactions from **A** positively regulated by oncogenic Kras (higher in 3w ON compared to 3d OFF) (adjusted p value <0.05). **C,** Circos plot showing average expression of fibroblast ligands connected to their predicted receptors on various cell populations as measured in the pancreatic single-cell RNA sequencing analysis. Ligands shown are from **B** (adjusted p value <0.05). **D,** Heatmap showing averaged single-cell RNA sequencing expression data (relative to the highest expressor) for genes in fibroblasts from a curated list of immunomodulatory factors. **E,** Violin plots showing expression of *Il33*, *Cxcl1, Il6*, *Saa3* and *Saa1* and their respective receptors *Il1rl1*, *Cxcr2*, *Il6ra*, *P2rx7* and *Scarb1* across all identified cell populations in both Kras* ON and OFF samples combined. See also Figure S4

To validate the mechanism of fibroblast reprogramming, we used an *in vitro* approach. We harvested conditioned media from iKras* cells derived from a Ptf1a^Cre/+^;R26^rtTa-ires-EGFP/rtTa-ires-EGFP^;TetO- Kras^G12D^;P53^R172H/+^ (iKras*P53*) mouse tumor (49). In brief, iKras*P53* cells (49) were grown in presence of DOX. Then, media was changed to either DOX-containing or DOX-free media, thus maintaining Kras* expression or inactivating it, respectively. After 48-72 hours, we harvested CM from each condition and boiled half of it to denature heat-labile components. CM was then administered to cultured primary mouse fibroblasts (CD1WT and B6318) derived from healthy adult pancreata as previously described (50) (44)) (**Fig. 5A**). Compared to regular media (control), CM from iKras* cells in presence of DOX induced expression of *Cxcl1, Il6, Il33* and *Saa3* in fibroblasts (**Fig. 5B and S5A**), while CM from iKras* cells grown without DOX failed to induce a similar level of expression. To distinguish between a Kras*-dependent protein factor or metabolite signaling to fibroblasts, we compared intact CM with boiled CM, and found that the latter induced *Il6* but none of the genes encoding the cytokines of interest (**Fig. 5B and S5A**). Thus, heat-labile factors such as proteins are likely involved in activating inflammatory gene expression, though a cancer cell-derived, Kras*-dependent metabolite may be responsible for *Il6* induction. Oncogenic Kras regulates the intracellular metabolome of pancreatic cancer cells (34). We enquired whether the extracellular metabolome is similarly dependent on expression of oncogenic Kras. Indeed, mass spectrometry-based metabolomics profiling revealed numerous extracellular metabolites whose abundance changed depending on Kras* expression (**Fig. S5B and S5C**). Among these, several released metabolites have been reported to act as direct extracellular signaling molecules (e.g. pyruvate, lactate, alpha-ketoglutarate) (**Fig. S5D**), while, alternatively, the decreased abundance of other metabolites may impact signaling downstream of mTOR (e.g. arginine, isoleucine, cystine, glutamine) (**Fig. S5C**) (51–54). We then interrogated the extracellular metabolite composition following incubation of the iKras* conditioned medium from the various conditions described above. We observed that the lactate concentration was not affected while pyruvate was depleted upon culturing with fibroblasts. Thus, it appears that fibroblasts consume pyruvate from the medium, an intriguing finding given that fibroblasts serve as a source of pyruvate in breast cancer (55).

**Figure 5.**
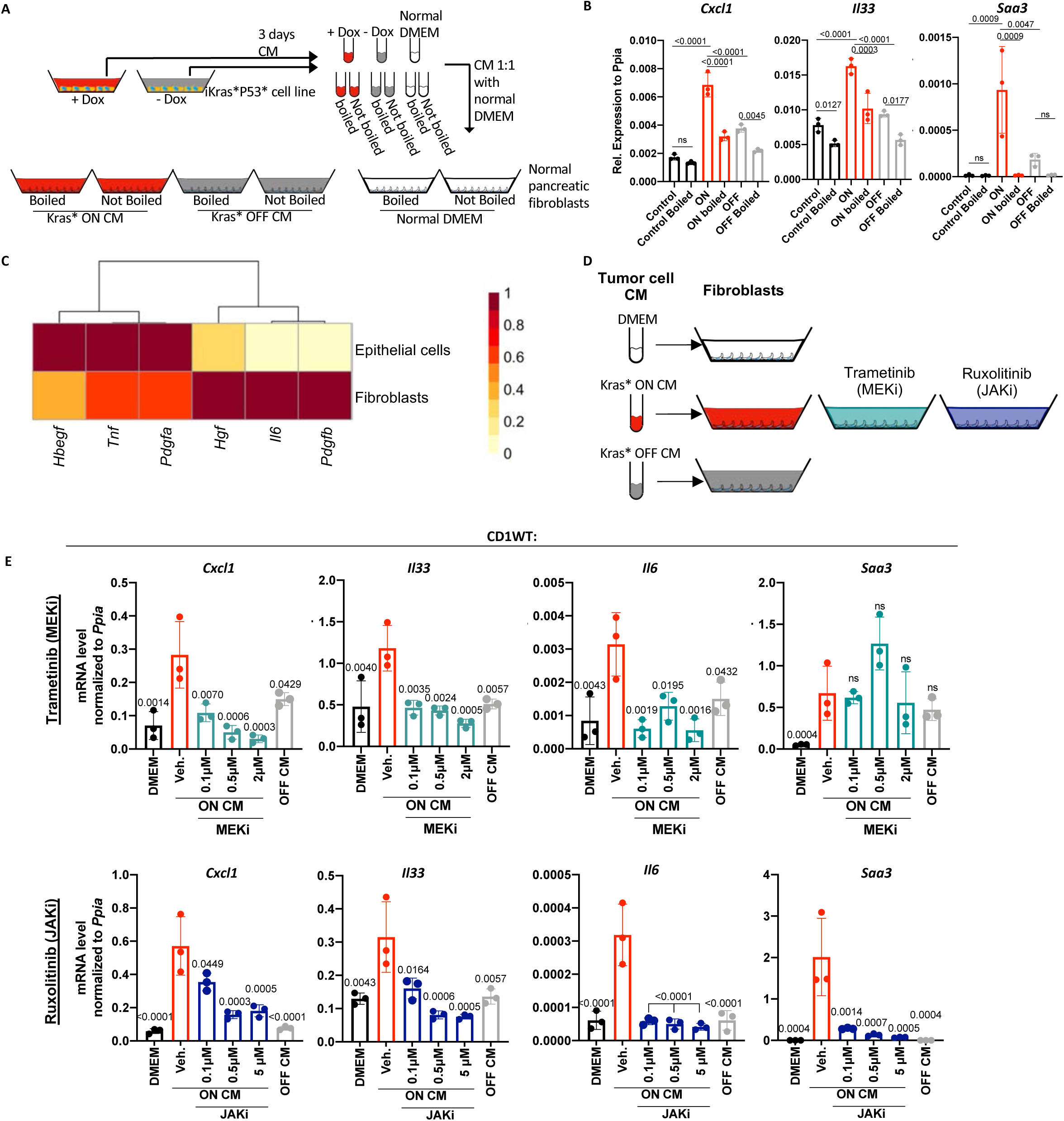
Fibroblasts are reprogrammed through Kras*-induced activation of the MAPK and JAK/STAT signaling pathways. **A,** Experimental design. Conditioned media (CM) was collected from iKras* cancer cells cultured with Doxycycline (+DOX) to activate Kras* expression (Kras* ON) or without Doxycycline (-DOX, Kras* OFF). The media samples were then boiled, to denature protein factors, or left intact, and used to culture pancreatic fibroblasts. DMEM was used as control. **B,** qRT-PCR for *Cxcl1, Il33* and *Saa3* expression in fibroblasts (CD1WT) that were cultured with CM either from Kras* ON, Kras* OFF or DMEM, either boiled or not boiled. Gene expression was normalized to *Ppia*. Data was shown as mean ± SD. N=3 per group. The statistic differences were determined by multiple comparison ANOVA. **C,** Heatmap showing averaged single-cell RNA sequencing expression data (relative to the highest expressor) for genes encoding for ligands that are secreted by fibroblasts and epithelial cells. **D,** Experimental design. Fibroblasts were exposed to iKras*P53* tumor cell CM either from Kras* ON, Kras* OFF or DMEM. Fibroblasts cultured with Kras* ON CM media were treated with either a MEK inhibitor (Trametinib), a JAK/STAT 2/3 inhibitor (Ruxolitinib) or vehicle. **E,** qRT-PCR results for *Cxcl1, Il33*, *Il6* and *Saa3* expression in fibroblasts (CD1WT) treated with CM as described in panel D. Data shown as mean ± SD. N=3 per group. The statistical differences were determined by multiple comparison ANOVA. See also Figure S5

We then interrogated the list of secreted factors expressed by transformed cells (likely within the CK19+ epithelial compartment) in the single cell sequencing data. We noted, among others, expression of *Hbegf, Tnf, Pdgf, Hgf* and *Il6*, while the corresponding receptors were expressed in fibroblasts (**Fig. 5C**). Expression of *Hbegf*, *Tnf* and *Pdgfa* was higher in epithelial cells, whereas *Hgf*, *Il6* and *Pdgfb* were higher in fibroblasts. Given the multitude of factors and their pleiotropic effect on intracellular pathway activation, we elected to block the common downstream effector pathways MAPK and JAK/STAT3 in fibroblasts rather than targeting individual secreted factors. We then repeated the experiment described above, this time inhibiting MAPK signaling (using Trametinib; MEKi) or JAK/STAT signaling (using Ruxolitinib; JAKi) (**Fig. 5D and S5E**). CM-induced fibroblast expression of *Cxcl1*, *Il33* and *Il6* was inhibited by both MEKi and JAKi, while *Saa3* expression was only inhibited by JAKi (**Fig. 5E** and **S5F**). Thus, tumor cell factors expressed upon Kras* activation reprogram fibroblasts by activating MAPK and JAK/STAT signaling to induce expression of a panel of inflammatory cytokines.

## DISCUSSION

Pancreatic cancer is associated with an extensive fibroinflammatory reaction that often constitutes over 80% of the tumor volume (for review see (8)). The interplay between tumor cells and components of the microenvironment promotes tumor growth and malignancy, and as such, is an area of intense investigation. What remains to be understood is how the stroma forms within the pancreas, as well as the events that lead from normal tissue to precursor lesions, and ultimately, malignant disease. Mutations in the KRAS gene are present in the majority of human pancreatic cancer instances (56). Further, those mutations occur early during carcinogenesis, and are prevalent in PanIN in the human pancreas (57). Interestingly, KRAS mutations occur spontaneously in a large proportion of individuals, increasing with age, yet pancreatic cancer remains a relatively rare disease (4, 5).

In the current study, we set out to understand how the interplay between epithelial cells expressing oncogenic Kras and the surrounding components of the pancreatic microenvironment regulate the balance between tissue repair and carcinogenesis. Using the iKras* genetically engineered mouse model (33) where oncogenic Kras can be activated and inactivated at will, we were able to follow changes in the pancreas at regular intervals immediately following the activation of oncogenic Kras. We found that Kras-dependent signals drive infiltration of monocyte-derived macrophages in the pancreas and activation of pancreatic fibroblasts. While changes in macrophages and fibroblasts appeared to occur simultaneously, we show that there is an epistatic relationship whereby inhibition of monocyte recruitment to the pancreas (by inhibition of CCR2) leads to reduced fibroblast activation, despite intact epithelial MAPK signaling. This finding also shows that recruitment of monocytes to the pancreas is important during the early stages of carcinogenesis, even as resident macrophages are equally required for lesion formation (38).

Given that myeloid cells are abundant during the early stages of carcinogenesis while T cell infiltration is rare, we focused the scope of this study on understanding epithelial cell/myeloid cell crosstalk. For this purpose, we let mice develop early pancreatic lesions and then inactivated oncogenic Kras. Our group has previously shown that inactivation of oncogenic Kras during these early stages of carcinogenesis leads to tissue repair over the course of two weeks, a process that requires macrophages (29, 33). We first hypothesized that continuous oncogenic Kras expression was necessary to maintain myeloid/macrophage infiltration in the pancreas. Contrary to our expectations, inactivation of oncogenic Kras had no effect on the number of myeloid cells present in the pancreas. However, analysis of the phenotype of infiltrating myeloid cells revealed a profound change in polarization. In the presence of oncogenic Kras, infiltrating macrophages express a series of markers that are consistent with tumor associated macrophages (TAMs), indicating that polarization of tumor-promoting macrophages occurs relatively early, prior to the formation of malignant disease. For example, macrophages in the neoplastic pancreas express the enzyme Arginase 1 (Arg1) and CD206/MRC1, while expression of iNOS, a marker of proinflammatory macrophages, is low. However, inactivation of oncogenic Kras leads to a reduction in Arg1^+^ macrophages.

To more comprehensively depict the relationship between epithelial and myeloid cells in early pancreatic tumorigenesis, we performed single cell RNA sequencing (scRNAseq) of iKras* mice either with active expression of oncogenic Kras or following 3 days of inactivation of oncogenic Kras. These timepoints were ideal as we have previously shown that at this interval, inactivation of oncogenic Kras causes levels of Kras-GTP (active Kras) to return to baseline while tissue repair is minimal, preventing us from comparing drastically different tissues (33). We then probed our scRNAseq dataset for known ligand-receptor pairs to determine which cell-cell signals were mediated by oncogenic Kras and responsible for macrophage re-polarization. Interestingly, most of the signals that changed upon inactivation of oncogenic Kras were derived from fibroblasts, rather than epithelial cells. Fibroblasts were a source of multiple cytokines: some were also expressed by other cell types (*Il33* and *Il6*), while others were uniquely fibroblast-specific (*Cxcl1, Saa1* and *Saa3*). Recently, IL33 has been described as expressed by epithelial cells during the early stages of carcinogenesis and noted to promote neoplastic progression (24); interestingly, we found that IL33 is also highly expressed by fibroblasts in early tumorigenesis stages, while its receptor is on multiple immune cell populations. Fibroblasts are also a source of IL6, with a known role in pancreatic cancer progression (58–60); Saa3 (ortholog to human SAA1), an apolipoprotein required for neoplastic progression (61) and CXCL1, which has been studied in epithelial cells, and shown to be a key moderator of T cell exclusion in malignant disease (62). In our study, fibroblasts emerged as mediators of epithelial/immune cell crosstalk during the onset of pancreatic carcinogenesis. To mechanistically dissect the fibroblast reprogramming process, we cultured primary mouse pancreatic fibroblasts with conditioned media from iKras*P53* tumor cells (49) expressing Kras* ON or OFF. We found that even in culture, fibroblasts are reprogrammed when exposed to CM from oncogenic Kras-expressing epithelial cells. Interestingly, while the majority of cytokines we assessed in fibroblasts required a heat labile component of the conditioned media, *Il6* did not, implying that a combination of Kras*-dependent signals are required for establishing a complete reprogramming of pancreatic fibroblasts, potentially including both peptide and metabolic secreted factors. Elucidating further details on these tumor-derived signals merits further exploration.

The role and origin of fibroblasts in pancreatic cancer remains poorly understood. Fibroblasts have been found to secrete a number of cytokines that influence the immune milieu (for example, CXCL12, which has been shown to reduce T cell infiltration in pancreatic cancer), and as such they promote cancer growth (63). However, in other models, fibroblast depletion promotes carcinogenesis (64). These contradictory findings might be explained by the heterogeneity of fibroblast populations. In recent years, the nature and origin of fibroblasts in advanced disease have been addressed by several studies, and previous assumptions regarding these cell populations have been challenged as a consequence. Cancer-associated fibroblasts (CAFs) have been long assumed to derive from a pancreatic stellate cell (PSC) population in the pancreas (similar to hepatic stellate cells), but recent lineage tracing work by our group has revealed that a substantial proportion of CAFs derive from perivascular fibroblasts present in the normal pancreas (65). Accordingly, a lineage tracing study following PSCs in pancreatic cancer showed that they only contribute to a small subset of CAFs (66). *In vitro* characterization and, more recently, scRNAseq studies have identified CAF subsets with specific transcriptional signatures and functional roles (67–69). Less studied is the role of fibroblasts in the neoplastic pancreas; our current results show that neoplasia-associated fibroblasts, reprogrammed by epithelial cells expressing oncogenic Kras, shape the immune microenvironment during the onset of carcinogenesis, likely initializing the immune suppressive nature of pancreatic neoplasia (25) (See Scheme in **Fig. 6**).

**Figure 6.**
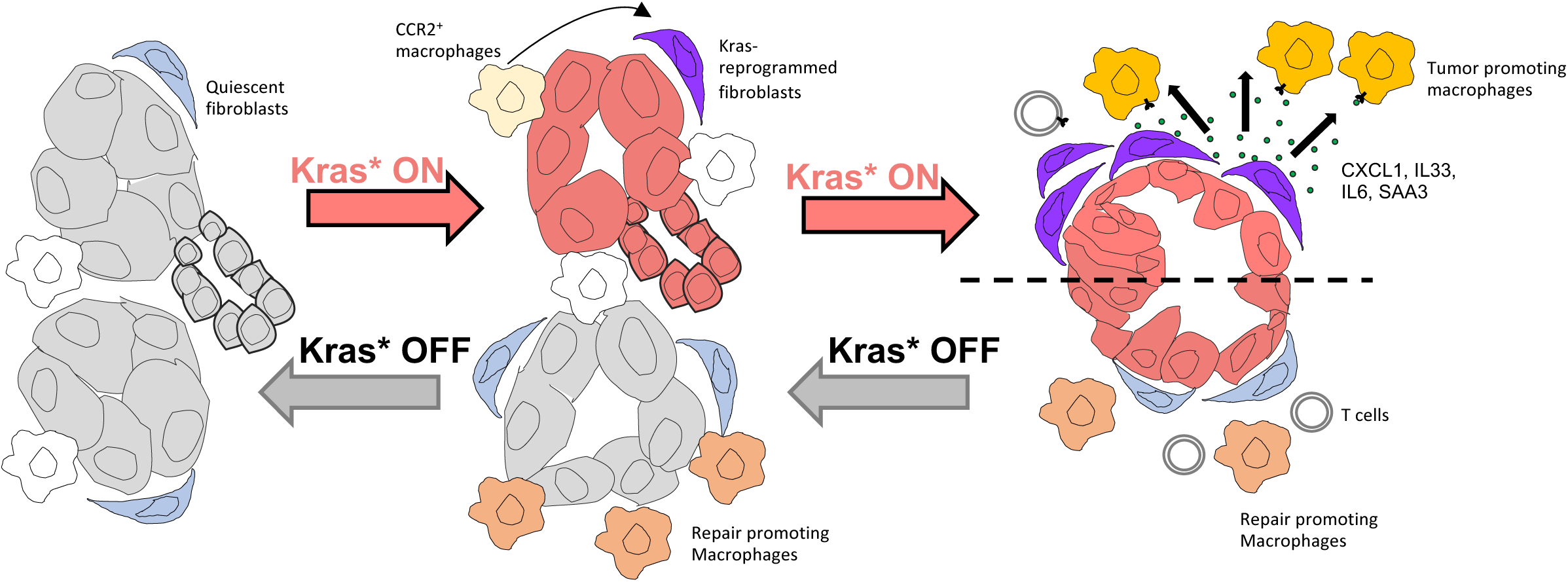
Working model. Upon Kras* expression, CCR2^+^ monocyte-derived macrophages infiltrate the pancreas, to complement pre-existing resident macrophages. Infiltrating CCR2^+^ macrophages are required to activate pancreatic fibroblasts. MAPK and JAK/STAT signaling are activated in fibroblasts at the PanIN stage, and induce expression of *Cxcl1, Il33, Il6,* and *Saa3.* These inflammatory molecules have receptors on immune cells, many of which are myeloid cells. In turn, the JAK/STAT3 pathway becomes activated in macrophages resulting in a tumor-promoting phenotype. Upon Kras* inactivation in the epithelial cells, inflammatory molecule expression is downregulated in fibroblasts, and macrophages are reprogrammed to a tissue repair phenotype that promotes the re-differentiation of acinar cells.

Recently, oncogenic Kras expression in pancreatic cancer has been associated with immune evasion (70). However, it remains to be seen whether the pancreatic cancer-associated immune response maintains its dependance on fibroblasts during late stages of carcinogenesis. In lung cancer, KRAS and MYC cooperate to shape the immune microenvironment (71); whether fibroblasts are required as signaling mediators in Kras-driven lung cancer and other diseases is not known.

In this study, we close the loop on communication between epithelial cells, fibroblasts and immune cells. Reprogramming of myeloid cells to alleviate the profound immune suppression of pancreatic cancer is a key concept in potential therapeutic approaches (72, 73). Unfortunately, the initial results from clinical trial testing of a CD40 agonist administered to reprogram myeloid cells combined with immune checkpoint inhibition and chemotherapy were disappointing (74), pointing to the need for yet additional avenues by which to target the microenvironment. Our current work suggests that fibroblast reprogramming should be considered in high-risk patients to prevent malignant progression and could also be explored to prevent tumor relapse in surgical patients. Finally, our study provides another piece of the puzzle in our understanding of the events that lead to the onset and progression of pancreatic cancer, and as such contribute to our fundamental knowledge of this deadly disease.

## ACKNOWLEDGEMENT

We thank Michael Scales and Dr. Benjamin L. Allen for providing the B6318 mouse wild-type fibroblast cell line. We thank Dr. Beth Moore for providing us with the CCR2 inhibitor. We thank Daniel Long for his histological services. This project was supported by NIH/NCI grants R01CA151588, R01CA198074, the University of Michigan Cancer Center Support Grant (NCI P30CA046592), the American Cancer Society to MPM and U01CA-224145 to MPM and HCC. YZ was funded by NCI-R50CA232985. AVD was supported by Rackham Merit Fellowship, Cellular Biotechnology Training Program (T32GM008353) and the NCI F31-CA247037. KLD, NGS and VRS were funded by the Cancer Biology Training Program T32-CA009676. NGS is also a recipient of the American Cancer Society Postdoctoral Award PF-19-096-01 and the Michigan Institute for Clinical and Healthy Research (MICHR) Postdoctoral Translational Scholar Program fellowship award. REM was supported by the NIH Cellular and Molecular Biology Training Grant T32-GM007315 and the Center for Organogenesis Training Program (NIH T32 HD007505). S.T & A.R were supported by institutional startup funds from the University of Michigan, NCI grants R37CA214955 & NCI P30CA046592 and a Research Scholar Grant from the American Cancer Society (RSG-16-005-01). SBK was supported by NIH T32-GM113900 and NCI F31-CA247076. SAK was supported by F31CA24745701. CAL was supported by the NCI (R37CA237421, R01CA248160) and UMCCC Core Grant (P30CA046592). FB was funded by the Association of Academic Surgery Joel Roslyn Award. TF was supported by (K08CA201581). Metabolomics studies performed at the University of Michigan were supported by NIH grant DK097153, the Charles Woodson Research Fund, and the UM Pediatric Brain Tumor Initiative. This project was also supported by the Tissue and Molecular Pathology and Flow Cytometry Shared Resources at the Rogel Cancer Center and the University of Michigan DNA Sequencing Core. CyTOF was performed at the University of Rochester University of Rochester Medical Center Flow Cytometry Shared Resource and at the Indiana University Simon Cancer Center Flow Cytometry Service.

## AUTHOR CONTRIBUTIONS

MPM directed the study. AVD, YZ and MPM designed experiments. AVD, KLD, KLB, WD, VIN, REM, ELLO, JL, WY, SBK, MB, FL, CAL, HCH, TLF performed the experiments and generated data. AVD, KLD, WD, NGS, ST, JL, VRS, SAK, ACN, FL, AR, FB, YZ and MPM analyzed and interpreted data. AVD, YZ and MPM wrote the manuscript, then all authors edited and approved the final version.

## METHODS

### Mice

Mice were housed in the specific pathogen-free animal facility at the Rogel Cancer Center, University of Michigan, and overseen by the unit for laboratory animal medicine (ULAM). Ptf1a(p48)- Cre;Rosa26^rtTa/rtTa^ mice were crossed to TRE-Kras^G12D^;R26^rtTa/rtTa^ to generate the p48-Cre;TRE- Kras^G12D^;R26^rtTa/rtTa^ (iKras*), as described (33). Expression of Kras* was induced in adult mice (8-14 weeks old) by replacing regular chow with Doxycycline chow (Bio-Serv, 1gm/kg). Acute pancreatitis was induced by administering eight hourly doses of caerulein (75μg/kg, Sigma-Aldrich) intraperitoneally for two consecutive days. For CCR2 inhibition, PF-04178903 (Pfizer, Inc, provided by Dr. Beth Moore) (Gurczynski, 2019, 30498200) was administered subcutaneously twice a day for 7 days at 50ug per dose in 100μl of PBS. PBS was used as control. All animal studies were conducted in compliance with the guidelines of the Institutional Animal Care & Use Committee (IACUC) at the University of Michigan.

### Histology and Immunohistochemistry

Pancreatic tissue samples from experimental and control mice were fixed in 10% neutral-buffered formalin (FisherBrand) overnight and then embedded in paraffin and sectioned into slides. Hematoxylin and Eosin (H&E), Gomori’s Trichome, Periodic Acid-Schiff (PAS) and Immunofluorescence (IF) staining were performed as previously described (33). For immunohistochemistry, fresh cut paraffin sections were re-hydrated using 2 series of xylene, 2 series of 100% ethanol and then 2 series of 95% ethanol. Water was used to wash all residues from previous washes. Antigen retrieval was performed using Antigen Retrieval CITRA Plus (BioGenex) and microwaved for total 8 minutes. Upon cool down, sections were blocked using 1% BSA in PBS for 30 minutes and then primary antibodies (for details see Supplementary Table 1) were used at their corresponding dilutions. Primary antibody incubation was performed at 4°C overnight. Biotinylated secondary antibodies were used in 1:300 dilution and applied to sections for 45 minutes at room temperature. Following secondary antibody incubation, sections were incubated for 30 minutes with the ABC reagent from Vectastain Elite ABC Kit (Peroxidase), followed by DAB (Vector). For immunofluorescence (IF), Alexa fluor secondary antibodies (Invitrogen) were used, then slides were mounted with Prolong Diamond Antifade Mountant with DAPI (Invitrogen). TSA Plus Fluorescein system (PerkinElmer) was used in IF for mouse primary antibodies. Olympus BX53F microscope, Olympus DP80 digital camera, and CellSens Standard software were used for imaging. Quantification of positive cell number or area was done by ImageJ using 3-5 images/slide (200x or 400x magnification) taken from 2-4 samples per group.

### Multiplex IHC Staining and Analysis

Multiplex immunohistochemistry staining was performed on paraffin embedded pancreatic tissue sections as follows. Slides were baked in a hybridization oven for one hour at 60 degrees Celsius, cooled for 10 minutes at room temperature, then dipped sequentially (x3) into xylene for 10 minutes each for removal of paraffin. Slides were then rehydrated in alcohol with dilutions of 100%, 95%, then 70% for 10 minutes each, followed by a wash in deionized water for 2 minutes. Slides were then placed in neutral buffered formalin for 30 minutes. The slides were then washed for 2 minutes in deionized water then microwaved at 100% power in Rodent Decloaker (Biocare Medical) for 30 seconds, the power level was reduced to 20% and microwaving continued for an additional 10 minutes followed by a resting step of 15 minutes at room temperature. Microwaving continued at 10% power for an additional 10 minutes. Prior to microwaving with Rodent Decloaker, plastic wrap was secured on top of the microwave-proof slide box with rubber bands and a partial opening for steam escape to prevent loss of solution. After the last microwaving step, slides were left to cool until slides and solution achieved room temperature. The multiplex staining was performed for each primary-color combination. Sides were placed in a deionized water wash for two minutes followed by a TBST wash for 2 minutes. Slides were placed in a slide incubation chamber and Bloxall was applied for 10 minutes followed by an additional blocking step of 1% BSA (in TBST) for 20 minutes, primary antibody was applied after tapping slide to remove the primary antibody and was left to incubate for 1 hour, slides were washed in TBST (x3) for 2 minutes each, secondary antibody was applied, followed by TBST wash (x3) for 2 minutes each, Opal color was applied for 10 minutes and a TBST wash (x3) for 2 minutes each was performed. The slides were then microwaved with either AR6 or AR9 for 45 seconds at 100% followed by 15 minutes at 20%. The previous steps were then repeated for each of the following antibodies and Opal colors in exact listed order: F480 at 1:600 (abcam ab6640), CD3 at 1:400 (Dako A0452)-TSA 520, CD8 at 1:400 (Cell Signaling 98941), Arg1 at 1:100 (Cell Signaling 93668), CK19 at 1:400 (Max Plank Institute Troma III). After the last application of multiplex was completed, sides were washed as above and placed in AR6, then microwaved. After cooling the slides were washed in deionized water followed by TBST for 2 minutes each. Opal spectral DAPI solution was applied (3 drops diluted in 1mL of TBST for 10 minutes followed by a wash in TBST for 30 seconds. Coverslips were mounted with Prolong Diamond, slides were left to lie flat overnight away from light. If the entire multiplex could completed without interruption, the slides were left in AR6 or AR9 after a microwaving step, covered from light until the next day. All primary antibodies were diluted in 1% BSA and all TSA Opal colors were diluted in TSA diluent at 1:50.

The Mantra Quantitative Pathology Work Station was used to image sections of each of the slides. One to three images per slide was captured at ×20 magnification. Cube filters (DAPI, CY3, CY5, CY7, Texas Red, Qdot) were used in taking each image capture. All images were analyzed using inForm Cell Analysis software (Akoya Biosciences). Sixty-three images of encompassing the 26 different control and experimental slides were batched analyzed after using a mouse formulated library consisting of each single TSA fluorophore listed above. Unmixing was performed and no spectral overlap was found between fluorophores. Using inForm version 2.3.0 training software, cell compartments were segmented into nucleus, cytoplasm, and membrane. DAPI was used to identify the nucleus of the cells and determine their shape and size. The cytoplasm was segmented using Arg1 (Opal 650), F480 (Opal 540), and CD3 (Opal 520). The inner distance to the nucleus was set at zero pixels and the outer distance of the nucleus was set at 6 pixels. The membrane was segmented using CK19 (Opal 690) with the max cell size set at 25 pixels from nucleus to the membrane, each pixel is 0.496 microns. X and Y coordinates were assigned to each identified cell in each image. Cell phenotypes (Tcell, macrophage, epithelial cell, acinar cell, other cell) were manually selected at random throughout the 63 images. Tcell, macrophage, and epithelial cells were selected based on single staining of CD3 (Opal 520), F480 (Opal 540), and CK19 (Opal 690) respectfully. Other cells were manually selected based on the lack of staining of the above listed fluorophores and acinar cells were labeled based on the lack of listed fluorophores and tissue morphology. Fluorescent intensity of Arg1 (Opal 650) in the nucleus at a max threshold of 200 and a positivity threshold level of 13 was used to identify Arg1^+^ cells. R programs were used to make complex phenotypes by combining the primary cell phenotypes (Tcell, macrophage, epithelial cell, acinar cell, and other cell) and the positivity thresholds for CD8+ and Arg1+.

### Flow cytometry

Pancreata were harvested and dissociated to single cells by mincing the tissue finely using scissors followed by Collagenase IV (1 mg/ml, Sigma) digestion for 30 minutes at 37°C while shaking. A 40um mesh strainer was used to separate single cells. RBC lysis buffer (eBioscience) was used to lyse all the red blood cells. Live cells were stained for surface markers using antibodies listed in Supplemental Table 2. The cell were either fixed after the primary antibody staining and use for analysis or the cells were fixed and permeabilized before intracellular staining using antibodies (Table S2). Flow-cytometric analysis was performed on the Cyan ADP analyzer (Beckman coulter) and the ZE5 analyzer (Bio-Rad). Data was analyzed using the FlowJo v10 software.

### CyTOF

Pancreas was harvested and disrupted to single cells as described above. Three mesh strainers were used: 500um, 100um and 40um, to separate single cells. Cells were washed twice in PBS and incubated with Cell-ID cisplatin (1.67 umol/L) for 5 min at room temperature (RT), as a viability marker. Surface and intracellular staining was performed as detailed in manufacturer instructions (Fluidigm) with the antibodies listed in Table 2. Cell were shipped in intercalator buffer on ice overnight to the Flow Cytometry core at the University of Rochester Medical Center, where sample preparation was finalized, and CyTOF2 Mass Cytometer (Helios) analysis was performed. Data analysis was performed using the Premium CytoBank Software (cytobank.org) and R studio using the cytof workflow from Nowicka M. Et. al. (75).

### Single-cell RNA sequencing

Pancreatic tissues were mechanically minced and enzymatically digested with collagenase IV (1 mg/ml in RPMI). Cell suspensions were subsequently filtered through 500-μm, 100-μm, and 40-μm mesh to obtain single cells. Dead cells were removed using the MACS Dead Cell Removal Kit (Miltenyi Biotec). The resulting single cell suspensions were pooled by experimental group (Kras ON/Kras OFF). Single-cell complementary DNA libraries were prepared and sequenced at the University of Michigan Advanced Genomics Core using the 10× Genomics Platform (Raw and processed data are available at GSM4175981 and GSE179846). Samples were run using 50-cycle paired-end reads on the HiSeq 4000 (Illumina) to a depth of 100,000 reads. The raw data were processed and aligned by the University of Michigan DNA Sequencing Core. Cell Ranger count version 3.0.0 was used with default settings, with an initial expected cell count of 10,000. R version 3.6.2, RStudio version 1.2.5033, and R package Seurat version 3.2.2 were used for scRNA-seq data analysis (RStudio Team RStudio: Integrated Development for R (RStudio, 2015); http://www.rstudio.com/; R Core Development Team R: A Language and Environment for Statistical Computing (R Foundation for Statistical Computing, 2017); https://www.R-project.org/ (76) (77) ). Data were initially filtered to only include cells with at least 100 genes and genes that appeared in more than three cells. Data were normalized using the NormalizeData function with a scale factor of 10,000 and the LogNormalize normalization method. Data were then manually filtered to include only cells with 1,000-60,000 transcripts and <15% mitochondrial genes. Variable genes were identified using the FindVariableFeatures function. Data were scaled and centered using linear regression of transcript counts. PCA was run with the RunPCA function using the previously defined variable genes. Cell clusters were identified via the FindNeighbors and FindClusters functions, using dimensions corresponding to approximately 90% variance as defined by PCA. UMAP clustering algorithms were performed with RunUMAP. Clusters were defined by user-defined criteria. The complete R script including figure-specific visualization methods is publicly available on GitHub (https://github.com/PascaDiMagliano-Lab/).

For Interactome analysis, ligand–receptor pairs were defined based on a curated literature-supported list in Ramilowski *et al.* (47), further curated as described in (48). The average expression values of ligands and receptors in each cell population for both experimental groups (Kras ON/Kras OFF) were calculated individually, and ligands and receptors expressed below a user-defined threshold (median average expression) were removed from analysis. Ligand–receptor pairs were also excluded if the ligands and receptors were not expressed in both experimental groups. Differences in the ligands and receptors between groups were determined using the Wilcoxon ranked test, and P values were adjusted for multiple comparisons using the Bonferroni correction method. Ligands and receptors were considered significantly different if P < 0.05. The resulting data table was visualized using Cytoscape (version 3.7.2) software (78). The complete R script is publicly available on GitHub (https://github.com/PascaDiMagliano-Lab/). The Circos plot was built with the Circos software (version 0.69-9) and the heatmap values within the circus plot displays the average expression of each fibroblast ligand and their corresponding receptor across the iKras* pancreatic tissue.

### Cell culture

All cell lines were cultured in DMEM with 10% Fetal Bovine Serum and 1% Penicillin Streptomycin. The tumor cell line (9805) was derived from an iKras*P53* (ptf1a-Cre; TetO-Kras^G12D^;Rosa26^rtTa/+;^ p53^R172H/+)^ pancreatic cancer (49). The fibroblasts cell lines, CD1WT and B6318, were derived from wild type pancreata of a mixed or C57BL/6J background, respectively. The 9805 cell line was maintained in Doxycycline-containing (1ug/mL) medium. To generate conditioned media, fresh medium containing Doxycycline (Kras* ON) or without Doxycycline (Kras* OFF) was replaced and harvested after 2-3 days based on cell confluency. The media were then centrifuged (300G, 10 minutes at 4°C) to remove contaminating cancer cells. The CM was used to culture fibroblasts with a 1:1 ratio of CM to normal DMEM for 2-3 days. An aliquot of the conditioned media before and after culturing fibroblasts was collected for metabolomics analysis (described below). To test the effect of heat labile factors, media were boiled for 10 mins to denature secondary and tertiary peptide structures. For inhibitor experiments, Ruxolitinib (INCB018424, Selleckchem) at 0.1 μM, 0.5 μM and 5 μM or Trametinib (GSK1120212, Selleckchem) 0.1 μM, 0.5 μM and 2 μM or vehicle control were added to the medium.

### Quantitative RT-PCR

RNA was extracted using the QIAGEN kit. RNA samples went through Reverse Transcription-PCR (RT-PCR) using the cDNA kit (Applied Biosystems). cDNA samples for quantitative Real Time PCR were prepared using a mix of 1X Fast-SYBR Green PCR Master Mix (Applied Biosystems) and the primers listed in Table 4. The reaction conditions were used as previously described (Collins et al. 2012). Cyclophilin A (*Ppia*) was used as housekeeping control.

### Western Blot

Cells were lysed using RIPA buffer (Sigma-Aldrich) with protease and phosphatase inhibitors (Sigma-Aldrich). Protein was quantified and the same amount of protein was loaded to the wells in a 4-15% SDS-PAGE gel (BioRad). Protein was transferred to a PVDF membrane (BioRad), that was blocked with milk and then incubated with primary antibodies listed in Supplementary Table 1 overnight. HRP-conjugated secondary anti-rabbit and anti-mouse (1:5000) were used and detected by using the enhance Chemiluminescent Substrate (PerkinElmer). The bands were visualized using the ChemiDoc Imaging System (BioRad).

### Metabolomics analysis

The conditioned medium from tumor cells and fibroblasts (described above) was aspirated and 1 ml of 80% methanol -cooled in dry ice-was added per well. The plates were incubated on dry ice for 10 min, then scraped and transferred to sample tubes. The samples were vortexed and centrifuged for 10 min at 13,000 x *g*, 4 °C. Then, 1 ml of the supernatant was aspirated from each tube, transferred to a tightly capped sample tube, and stored at -80 °C until analysis.

Samples were run on an Agilent 1290 Infinity II LC -6470 Triple Quadrupole (QqQ) tandem mass spectrometer (MS/MS) system with the following components: Agilent Technologies Triple Quad 6470 LC-MS/MS system with a 1290 Infinity II LC Flexible Pump (Quaternary Pump), Multisampler and Multicolumn Thermostat with 6 port valve. Agilent Masshunter Workstation Software LC/MS Data Acquisition for 6400 Series Triple Quadrupole MS with Version B.08.02 was used for compound optimization, calibration, and data acquisition.

The following solvents were employed for LC analysis. Solvent A (97% water and 3% methanol 15 mM acetic acid and 10 mM tributylamine, pH of 5), solvent B (15 mM acetic acid and 10 mM tributylamine in methanol) and washing solvent C (acetonitrile). LC system seal washing solvent (90% water and 10% isopropanol) and needle wash solvent (75% methanol, 25% water) were used. Solvents were purchased from the following vendors: GC-grade tributylamine 99% (ACROS ORGANICS), LC/MS grade acetic acid Optima (Fisher Chemical), InfinityLab deactivator additive, ESI –L Low concentration Tuning mix (Agilent Technologies), LC-MS grade water, acetonitrile, methanol (Millipore) and isopropanol (Fisher Chemical).

For LC analysis, 2 µl of sample was injected into an Agilent ZORBAX RRHD Extend-C18 column (2.1 × 150 mm, 1.8 µm) with ZORBAX Extend Fast Guards. The LC gradient profile was: flow rate: 0.25 ml/min, 0-2.5 min, 100% A; 2.5-7.5 min, 80% A and 20% B; 7.5-13 min 55% A and 45% B; 13-24 min, 1% A and 99% B; 24-27min, 1% A and 99% C; 27-27.5min, 1% A and 99% C. Then, at 0.8 ml/min, 27.5-31.5 min, 1% A and 99% C; at 0.6 ml/min, 31.5-32.25 min, 1% A and 99% C; at 0.4 ml/min, 32.25-39.9 min, 100% A; at 0.25 ml/min, 40 min, 100% A. Column temp was maintained at 35 °C while the samples were at 4 °C.

For MS analysis, a 6470 Triple Quad MS calibrated with the Agilent ESI-L low concentration tuning mix was used. Source parameters: Gas temp 150°C, gas flow 10 l/min, nebulizer 45 psi, sheath gas temp 325°C, sheath gas flow 12 l/min, capillary -2000 V, Delta EMV -200 V. Dynamic MRM scan type was used with 0.07 min peak width and 24 min acquisition time. Delta retention time of ± 1 min, fragmentor of 40 eV and cell accelerator of 5 eV were incorporated in the method.

The MassHunter Metabolomics Dynamic MRM Database and Method was used for target identification. Key parameters of AJS ESI were: Gas Temp: 150°C, Gas Flow 13 l/min, Nebulizer 45 psi, Sheath Gas Temp 325°C, Sheath Gas Flow 12 l/min, Capillary 2000 V, Nozzle 500 V. Detector Delta EMV(-) 200.The QqQ data were pre-processed with an Agilent MassHunter Workstation QqQ Quantitative Analysis Software (B0700). The abundance level of each metabolite in every sample was divided by the median of all abundance levels across all samples for proper comparisons, statistical analyses, and visualization. The statistical significance test was done by a two-tailed t-test with a significance threshold level of 0.05.

Heatmaps were generated with Morpheus, https://software.broadinstitute.org/morpheus.

### Statistics

We used GraphPad Prism Version 8 software for most of our analysis. The normality was checked in all data sets and either T-test or Mann Whitney was performed for statistical analysis, statistically significant when p<0.05. qRT-PCR data was analyzed using multiple comparison ANOVA and considered statistically significant when p<0.05.

