## Supplemental Figures for "Extrinsic KRAS signaling shapes the pancreatic microenvironment through fibroblast reprogramming"

### Slide 1
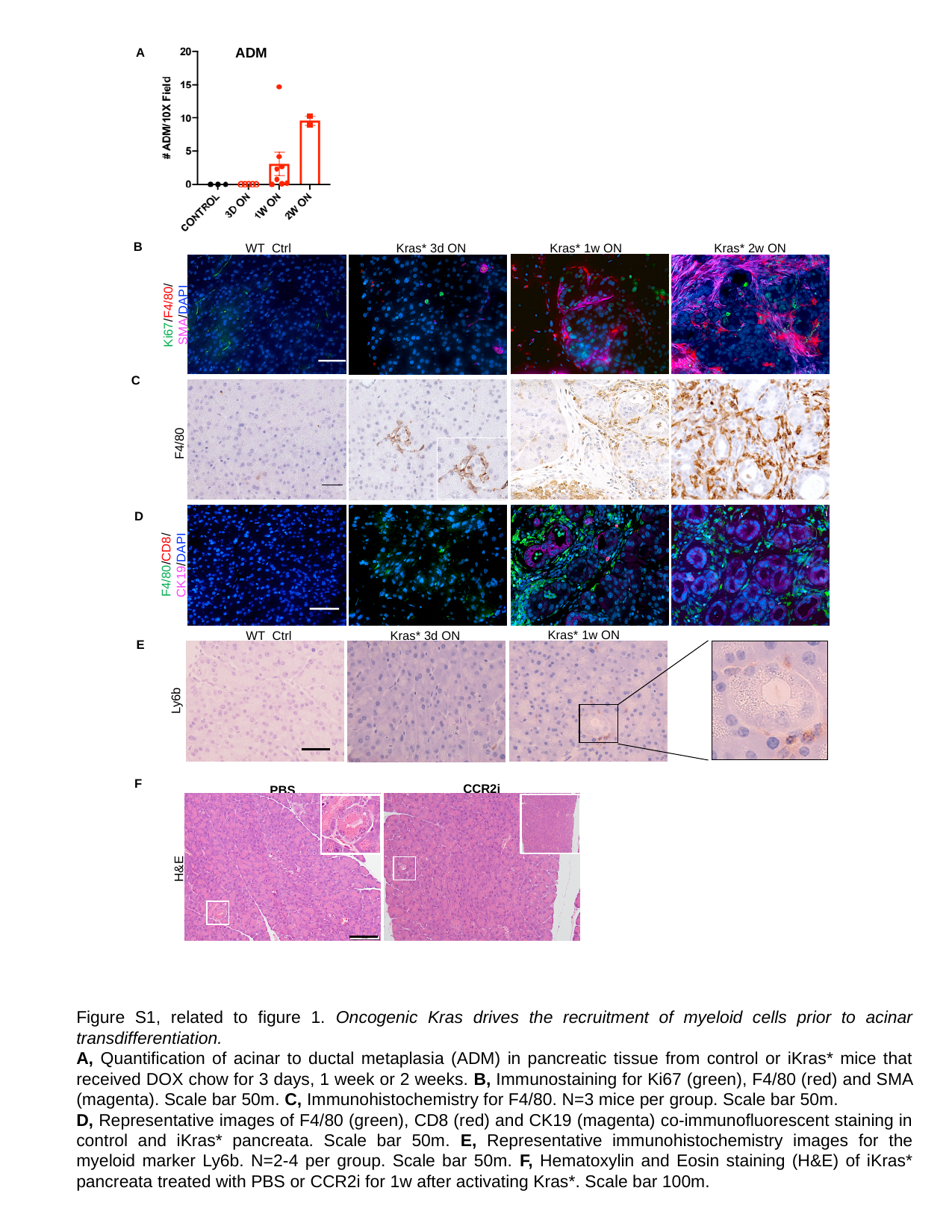

ADM
A
B
Ki67/F4/80/
SMA/DAPI
C
F4/80
D
F4/80/CD8/
CK19/DAPI
E
Kras* 1w ON
WT Ctrl
Kras* 3d ON
Ly6b
F
CCR2i
PBS
H&E
WT Ctrl
Kras* 1w ON
Kras* 2w ON
Kras* 3d ON

### Slide 2
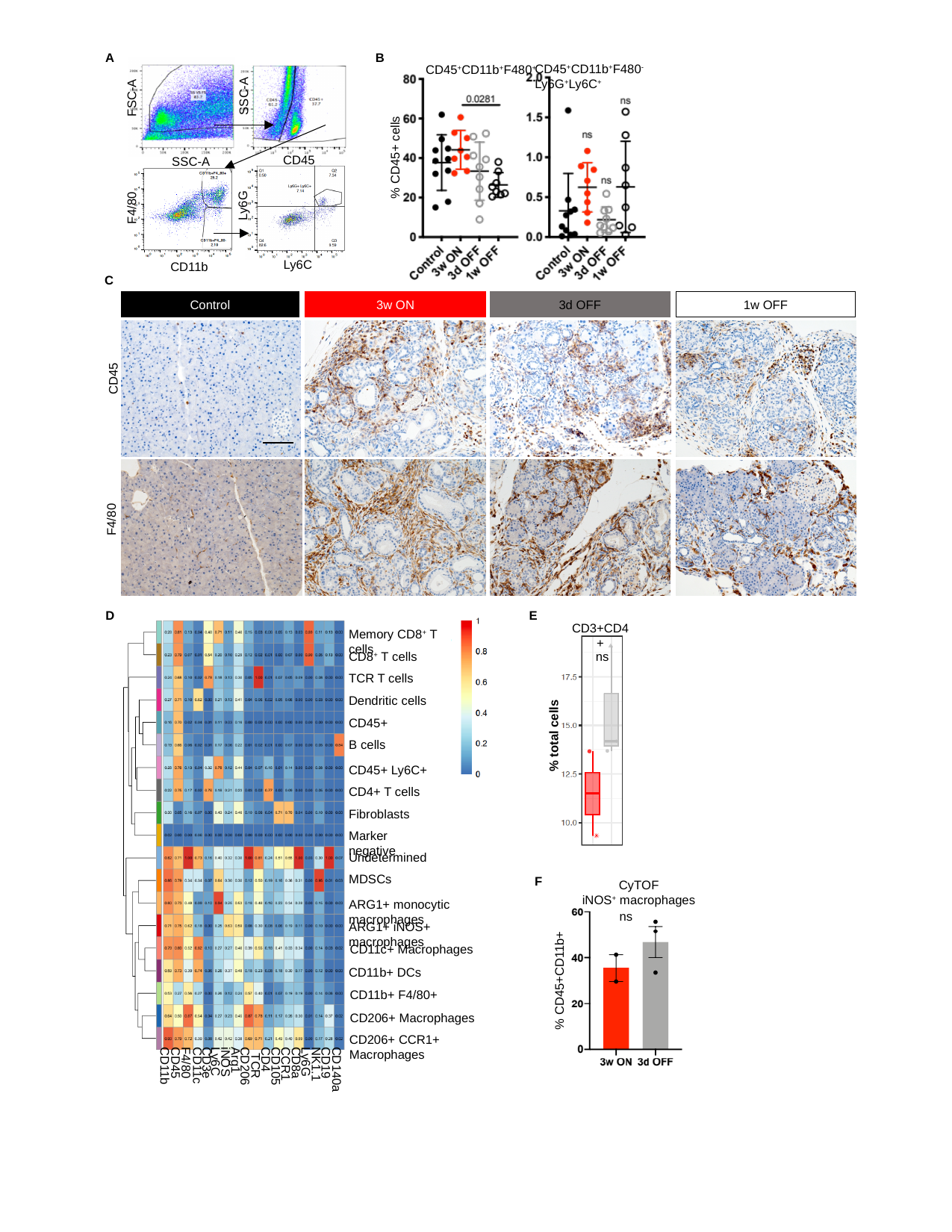

FSC-A
SSC-A
CD45
SSC-A
Ly6G
F4/80
Ly6C
CD11b
A
B
CD45^+^CD11b^+^F480^-^Ly6G^+^Ly6C^+^
CD45^+^CD11b^+^F480^+^
% CD45+ cells
C
Control
3w ON
3d OFF
1w OFF
CD45
F4/80
D
E
CD3+CD4+
% total cells
Memory CD8^+^ T cells
CD8^+^ T cells
Dendritic cells
CD45+
B cells
CD45+ Ly6C+
CD4+ T cells
Fibroblasts
Marker negative
Undetermined
MDSCs
ARG1+ monocytic macrophages
ARG1+ iNOS+ macrophages
CD11c+ Macrophages
CD11b+ DCs
CD11b+ F4/80+
CD206+ Macrophages
CD206+ CCR1+ Macrophages
ns
F
CyTOF
iNOS^+^ macrophages
% CD45+CD11b+
ns
CD4
Arg1
Ly6C
iNOS
Ly6G
CD45
CD3e
CD8a
CD19
F4/80
CCR1
NK1.1
CD11c
CD11b
CD206
CD105
CD140a

### Slide 3
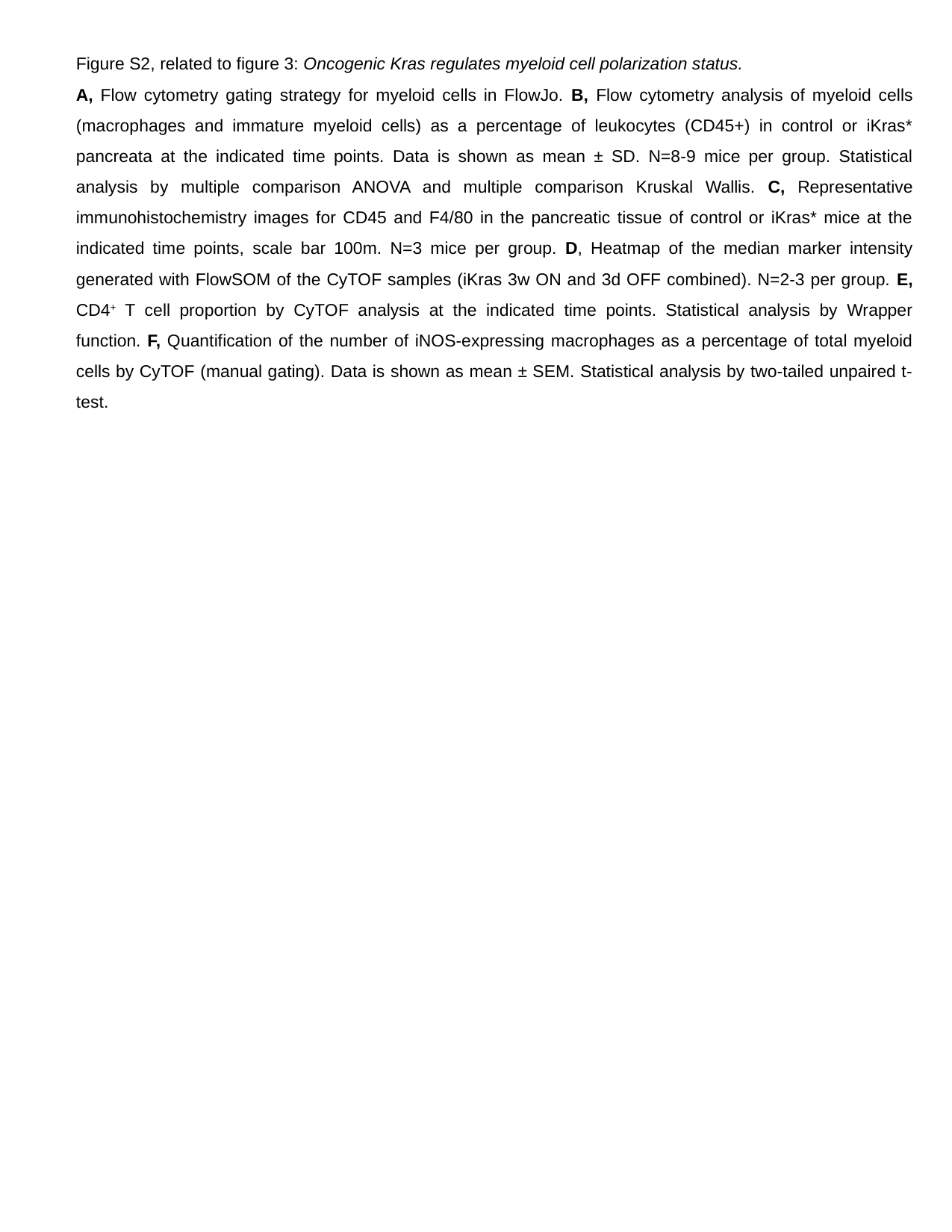

### Slide 4
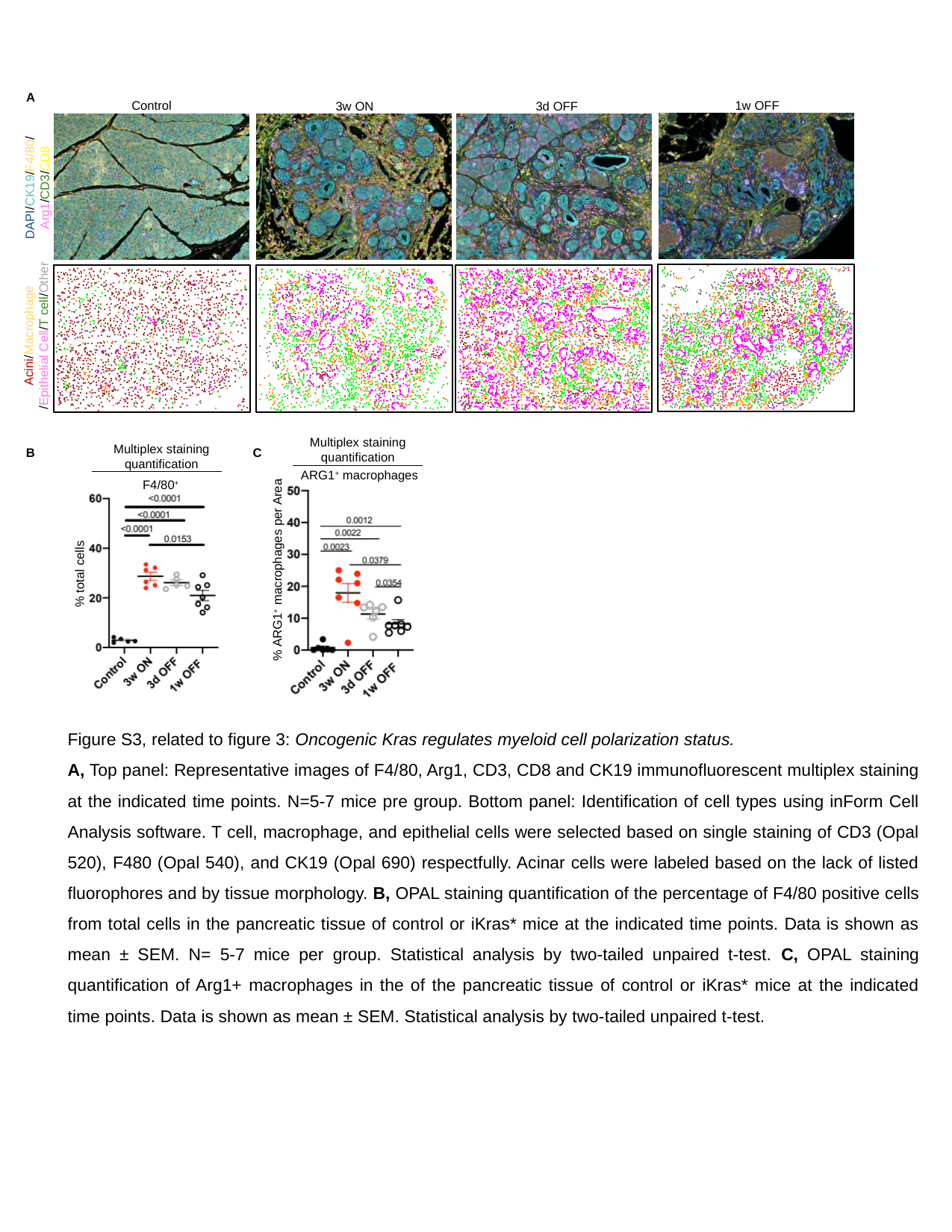

A
1w OFF
Control
3d OFF
3w ON
DAPI/CK19/F4/80/ Arg1/CD3/CD8
B
Acini/Macrophage
/Epithelial Cell/T cell/Other
Multiplex staining quantification
Multiplex staining quantification
C
ARG1^+^ macrophages
% ARG1^+^ macrophages per Area
F4/80^+^
% total cells
Figure S3, related to figure 3: Oncogenic Kras regulates myeloid cell polarization status.
A, Top panel: Representative images of F4/80, Arg1, CD3, CD8 and CK19 immunofluorescent multiplex staining at the indicated time points. N=5-7 mice pre group. Bottom panel: Identification of cell types using inForm Cell Analysis software. T cell, macrophage, and epithelial cells were selected based on single staining of CD3 (Opal 520), F480 (Opal 540), and CK19 (Opal 690) respectfully. Acinar cells were labeled based on the lack of listed fluorophores and by tissue morphology. B, OPAL staining quantification of the percentage of F4/80 positive cells from total cells in the pancreatic tissue of control or iKras* mice at the indicated time points. Data is shown as mean ± SEM. N= 5-7 mice per group. Statistical analysis by two-tailed unpaired t-test. C, OPAL staining quantification of Arg1+ macrophages in the of the pancreatic tissue of control or iKras* mice at the indicated time points. Data is shown as mean ± SEM. Statistical analysis by two-tailed unpaired t-test.

### Slide 5
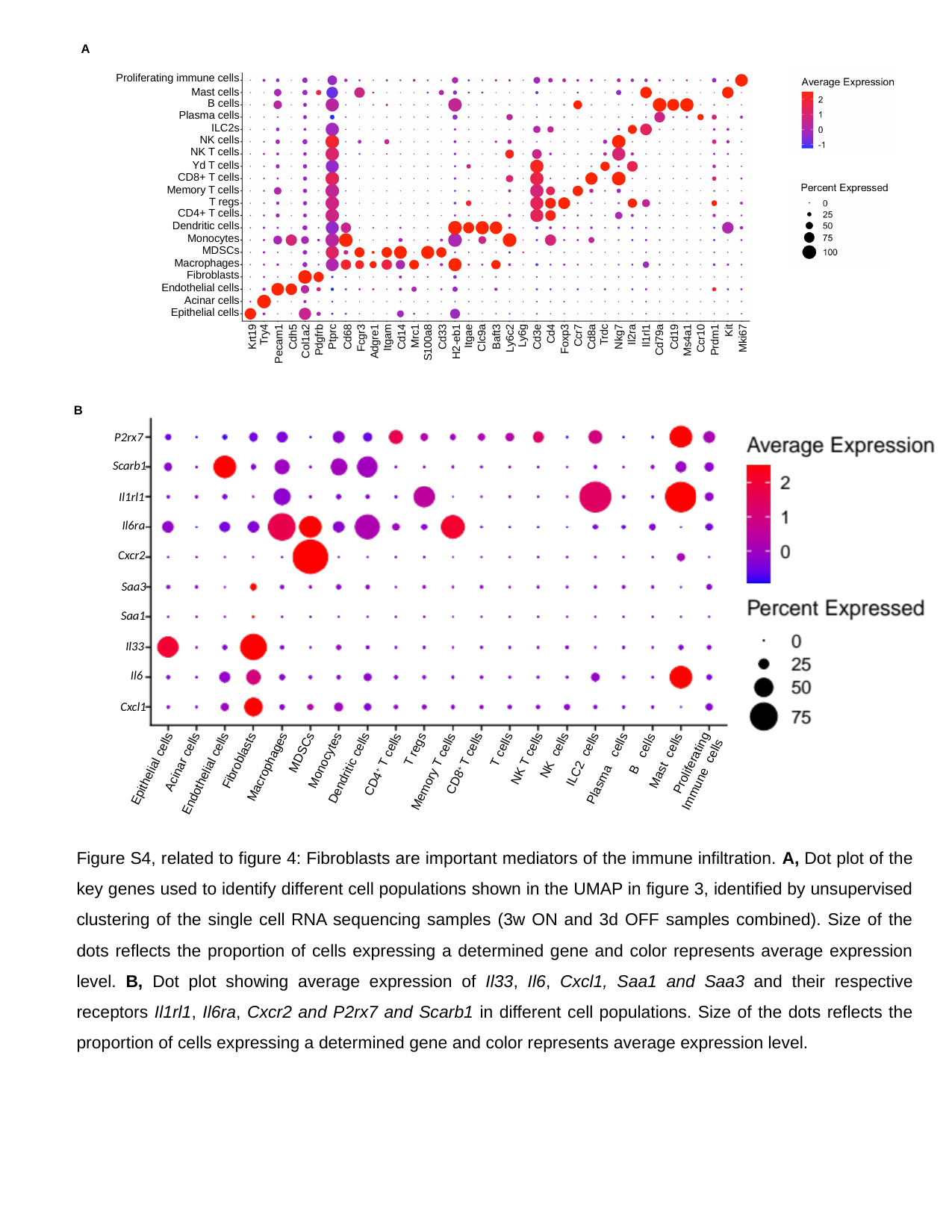

A
Proliferating immune cells
Mast cells
B cells
Plasma cells
ILC2s
NK cells
NK T cells
Yd T cells
CD8+ T cells
Memory T cells
T regs
CD4+ T cells
Dendritic cells
Monocytes
MDSCs
Macrophages
Fibroblasts
Endothelial cells
Acinar cells
Epithelial cells
Krt19
Try4
Pecam1
Cdh5
Col1a2
Pdgfrb
Ptprc
Cd68
Fcgr3
Adgre1
Itgam
Cd14
Mrc1
S100a8
Cd33
H2-eb1
Itgae
Clc9a
Baft3
Ly6c2
Ly6g
Cd3e
Cd4
Foxp3
Ccr7
Cd8a
Trdc
Nkg7
Il2ra
Il1rl1
Cd79a
Cd19
Ms4a1
Ccr10
Prdm1
Kit
Mki67
B
P2rx7
Scarb1
Il1rl1
Il6ra
Cxcr2
Saa3
Saa1
Il33
Il6
Cxcl1
Proliferating Immune cells
Endothelial cells
Acinar cells
Fibroblasts
Macrophages
MDSCs
Monocytes
Dendritic cells
CD8^+^ T cells
NK T cells
NK cells
ILC2 cells
Epithelial cells
Plasma cells
T regs
Memory T cells
CD4^+^ T cells
Mast cells
B cells
Figure S4, related to figure 4: Fibroblasts are important mediators of the immune infiltration. A, Dot plot of the key genes used to identify different cell populations shown in the UMAP in figure 3, identified by unsupervised clustering of the single cell RNA sequencing samples (3w ON and 3d OFF samples combined). Size of the dots reflects the proportion of cells expressing a determined gene and color represents average expression level. B, Dot plot showing average expression of Il33, Il6, Cxcl1, Saa1 and Saa3 and their respective receptors Il1rl1, Il6ra, Cxcr2 and P2rx7 and Scarb1 in different cell populations. Size of the dots reflects the proportion of cells expressing a determined gene and color represents average expression level.

### Slide 6
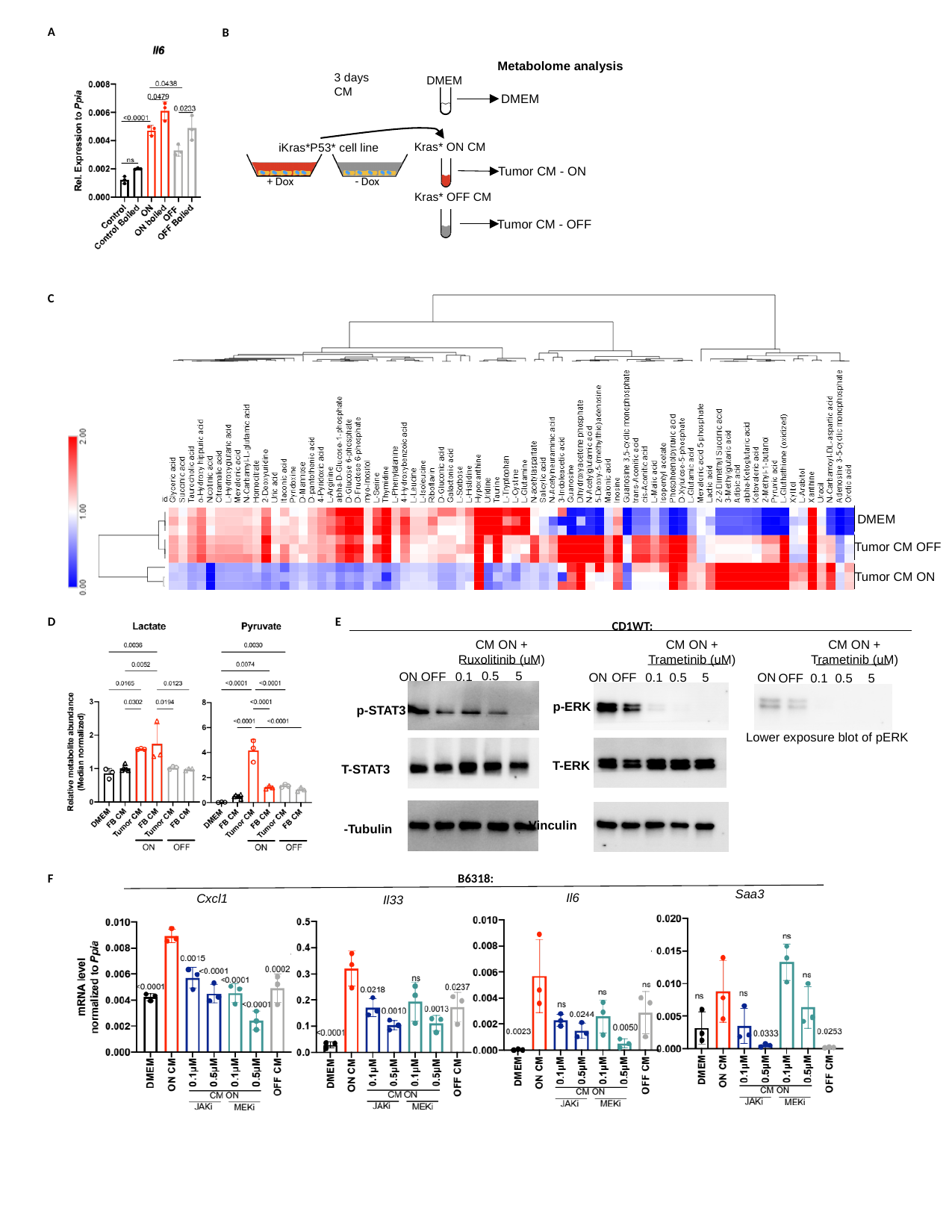

A
B
Metabolome analysis
3 days CM
DMEM
Kras* ON CM
Kras* OFF CM
iKras*P53* cell line
- Dox
+ Dox
DMEM
Tumor CM - ON
Tumor CM - OFF
C
DMEM
Tumor CM OFF
Tumor CM ON
E
F
D
CD1WT:
CM ON + Ruxolitinib (uM)
0.5
5
0.1
ON
OFF
CM ON + Trametinib (uM)
CM ON + Trametinib (uM)
0.1
0.5
5
ON
ON
OFF
OFF
0.5
0.1
5
p-ERK
p-STAT3
Lower exposure blot of pERK
T-ERK
T-STAT3
Vinculin
B6318:
Saa3
Il6
Cxcl1
Il33

### Slide 7
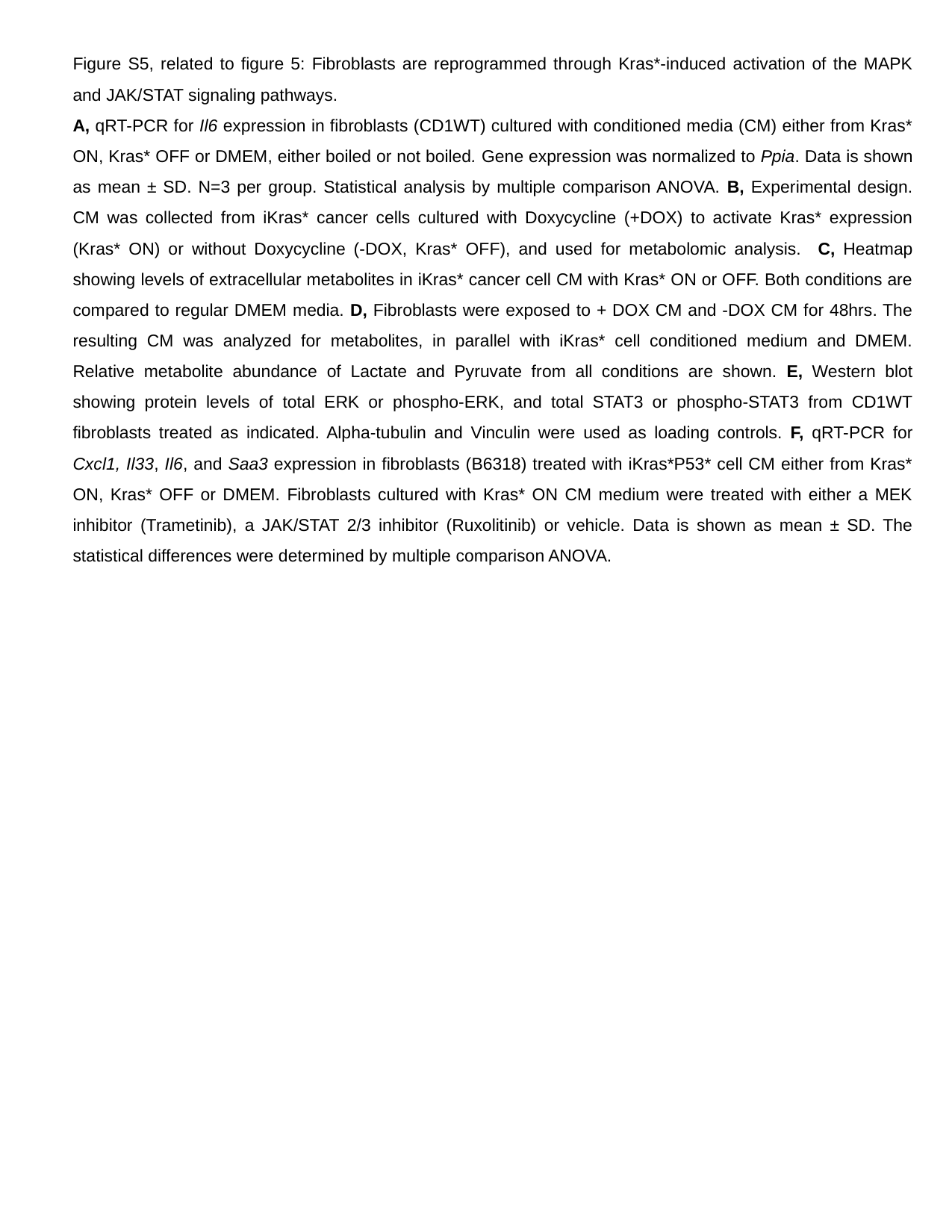

Figure S5, related to figure 5: Fibroblasts are reprogrammed through Kras*-induced activation of the MAPK and JAK/STAT signaling pathways.
A, qRT-PCR for Il6 expression in fibroblasts (CD1WT) cultured with conditioned media (CM) either from Kras* ON, Kras* OFF or DMEM, either boiled or not boiled. Gene expression was normalized to Ppia. Data is shown as mean ± SD. N=3 per group. Statistical analysis by multiple comparison ANOVA. B, Experimental design. CM was collected from iKras* cancer cells cultured with Doxycycline (+DOX) to activate Kras* expression (Kras* ON) or without Doxycycline (-DOX, Kras* OFF), and used for metabolomic analysis. C, Heatmap showing levels of extracellular metabolites in iKras* cancer cell CM with Kras* ON or OFF. Both conditions are compared to regular DMEM media. D, Fibroblasts were exposed to + DOX CM and -DOX CM for 48hrs. The resulting CM was analyzed for metabolites, in parallel with iKras* cell conditioned medium and DMEM. Relative metabolite abundance of Lactate and Pyruvate from all conditions are shown. E, Western blot showing protein levels of total ERK or phospho-ERK, and total STAT3 or phospho-STAT3 from CD1WT fibroblasts treated as indicated. Alpha-tubulin and Vinculin were used as loading controls. F, qRT-PCR for Cxcl1, Il33, Il6, and Saa3 expression in fibroblasts (B6318) treated with iKras*P53* cell CM either from Kras* ON, Kras* OFF or DMEM. Fibroblasts cultured with Kras* ON CM medium were treated with either a MEK inhibitor (Trametinib), a JAK/STAT 2/3 inhibitor (Ruxolitinib) or vehicle. Data is shown as mean ± SD. The statistical differences were determined by multiple comparison ANOVA.
