## Supplemental tables for "Extrinsic KRAS signaling shapes the pancreatic microenvironment through fibroblast reprogramming"

**Table S1:** IHC, IF and Western Blot antibodies

| Antibody | Supplier | Catalog # | IHC dilution | IF dilution | WB dilution |
| --- | --- | --- | --- | --- | --- |
| CD45 | BD Pharmingen | 553076 | 1:200 |  |  |
| F4/80 | Cell Signaling | 70076S | 1:250 | 1:250 |  |
| CD4 | Cell Signaling | 25229S | 1:00 |  |  |
| CD8 | Cell Signaling | 98941S | 1:400 | 1:100 |  |
| Foxp3 | Cell Signaling | 12653S | 1:100 |  |  |
| Arginase 1 | Cell Signaling | 93668S |  | 1:250 |  |
| aSMA | Sigma Aldrich | A2547 |  | 1:1000 |  |
| CK19 (Troma III) | Iowa Development Hybridoma Bark | - |  | 1:50 |  |
| Perk1/2 | Cell Signaling | 4370L |  | 1:100 |  |
| Ki67 | Abcam | ab15580 |  | 1:100 |  |
| CC3 | Cell Signaling | 9661L |  |  |  |
| E-cadherin | Cell Signaling | 14472S |  | 1:100 |  |
| CD3 | ABCAM | ab5690 |  | 1:100 |  |
| pStat3 Y705 | Cell Signaling | 9145S |  |  | 1:500 |
| Total STAT3 | Cell Signaling | 9139S |  |  | 1:1000 |
| p-ERK(T202/Y204) | Cell Signaling | 470L |  |  | 1:1000 |
| Total ERK | Cell Signaling | 4695S |  |  | 1:1000 |
| Vinculin | Cell Signaling | 13901 |  |  | 1:2000 |
| $a$-tubulin | Cell Signaling | 3873S |  |  | 1:2000 |

**Table S2:** Flow Cytometry antibodies

| Antibodies | Supplier | Catalog # | Clone | Dilution |
| --- | --- | --- | --- | --- |
| CD45 | Invitrogen | MCD4530 | 30-F11 | 1:100 |
| CD45 | BD Horizon | 563891 | 30-F11 | 1:100 |
| CD11b | BD Pharmingen | 557657 | M1/70 | 1:100 |
| F4/80 | Invitrogen | 15-4801-82 | BM8 | 1:100 |
| Ly6G | BD Pharmingen | 551460 | 1A8 | 1:100 |
| Ly6C | BD Pharmingen | 560592 | AL-21 | 1:100 |
| Arg1 | R&D | IC5868F | Polyclonal | 1:50 |
| iNOS | Invitrogen | 12-5920-82 | CXNFT | 1:100 |
| CD206 | BD Pharmingen | 565250 | MR5D3 | 1:100 |

**Table S3:** CyTOF antibodies

| Antibodies |  | Clone | Isotope | Dilution |
| --- | --- | --- | --- | --- |
| CD45 | Fluidigm | 30-F11 | Y89 | 1:200 |
| Ly6G | Fluidigm | 1AB | Pr141 | 1:400 |
| CD11b | Fluidigm | M1/70 | Nd143 | 1:300 |
| CD4 | Fluidigm | RM4/5 | Nd145 | 1:200 |
| F480 | Fluidigm | BM8 | Nd146 | 1:100 |
| CD140a | Fluidigm | APA5 | Nd148 | 1:100 |
| CD19 | Fluidigm | 6D5 | Nd149 | 1:200 |
| Ly6C | Fluidigm | HK1.4 | Nd150 | 1:400 |
| CD3e | Fluidigm | 145-2C11 | Sm152 | 1:100 |
| CD105 | Custom | MJ7/18 | Dy164 | 1:100 |
| TCR gd | Fluidigm | GL3 | Tb159 | 1:100 |
| CD191 CCR1 | Custom | S15040E | Gd160 | 1:100 |
| iNOS | Fluidigm | CXNFT | Dy161 | 1:100 |
| Arginase1 | Custom | Polyclonal | Er166 | 1:400 |
| CD8a | Fluidigm | 53-6.7 | Er168 | 1:200 |
| CD206 | Fluidigm | C068C2 | Tm169 | 1:200 |
| CD161 NK1.1 | Fluidigm | PK136 | Er170 | 1:100 |
| CD11c | Fluidigm | N418 | Bi209 | 1:100 |

**Table S4:** Primers for quantitative RT-PCR

| Genes | Forward Primer | Reverse Primer |
| --- | --- | --- |
| Cxcl1 | 5’ CTGGGATTCACCTCAAGAACATC 3’ | 5’ CAGGGTCAAGGCAAGCCTC 3’ |
| Il6 | 5’ TTCCATCCAGTTGCCTTCTTGG 3’ | 5’ TTCTCATTTCCACGATTTCCCAG 3’ |
| Il33 | 5’ TGAGACTCCGTTCTGGCCTC 3’ | 5’ CTCTTCATGCTTGGTACCCGA T 3’ |
| Saa3 | 5’ TGCCATCATTCTTTGCATCTTGA 3’ | 5’ CCGTGAACTTCTGAACACCCT 3’ |

**Table S5:** Multiplex IHC antibodies and cell phenotyping

| **Complex Phenotype** | **Opal** | **Primary Phenotype** | **Antibody Scoring** | **Cell segment** | **Mean signal intensity** |
| --- | --- | --- | --- | --- | --- |
| Macrophage | F4/80-Opal 540 | Macrophage | None | n/a | n/a |
| Arg1^+^ macrophage | Arg1-Opal 650 | Macrophage | Arg1 | Nucleus | 13 and above |
| Arg1^-^ macrophage | - | Macrophage | Arg1 | Nucleus | <13 |
| Epithelial cell | CK19-Opal 690 | Epithelial | None | n/a | n/a |
| Acinar cell | - | Acinar cell | None | n/a | n/a |
